# Admixture influences the genetic architecture of DNA methylation in a wild primate hybrid zone

**DOI:** 10.1101/2025.08.08.669436

**Authors:** Tauras P. Vilgalys, Jordan A. Anderson, Arielle S. Fogel, Dana Lin, Elizabeth A. Archie, Susan C. Alberts, Jenny Tung

## Abstract

**Background:** Hybrid zones play a central role in evolutionary biology because they serve as natural laboratories for studying how traits and taxa diverge. Changes in gene regulation make important contributions to this process. However, the degree to which admixture shapes gene regulatory variation in hybrid populations remains poorly understood. Here, we combine genome-wide resequencing and DNA methylation data from 295 hybrid baboons—members of a single, intensively studied natural population—to investigate how admixture affects the genetic architecture of this important epigenetic mark.

**Results:** We find that local genetic ancestry frequently predicts DNA methylation levels and recapitulates differences between the parent species. By performing methylation quantitative trait locus (meQTL) mapping, we show that these differences predominantly arise due to evolved differences in allele frequencies. Thus, admixture in the hybrid population increases variance in DNA methylation, including by introducing meQTL that would otherwise be invariant. Finally, we integrate massively parallel reporter assay data to show that admixture-derived variation in DNA methylation alters enhancer activity and gene expression.

**Conclusions:** Together, these results demonstrate how admixture can meaningfully alter the genetic architecture of gene regulatory traits in natural hybrid zones. They also suggest that the genetic architecture of DNA methylation is conserved across closely related primates, suggesting that genetic effects on gene regulation may remain stable over timescales that range into the millions of years.

## Background

Hybrid zones have long held a special place in evolutionary biology as “natural laboratories” for studying the evolutionary process, including the genetic basis of traits that have diverged between the parent species [1,2]. For example, in Darwin’s finches and European crows, naturally occurring hybrids have been leveraged to identify the genetic loci that underlie species-level differences in adaptively important traits like plumage color and beak size [3,4]. Natural hybridization can also introduce new, functionally important genetic variation. In humans, for instance, alleles introduced via admixture with archaic hominins have facilitated adaptation to high-altitude environments and resistance to pathogens [5–7], including through the re-introduction of alleles previously lost in our own lineage [8].

In many of these cases, the phenotypic effect of introgressed variation is thought to occur through a proximate effect on gene regulation [9]. These observations fit with experimental studies where crosses in captivity not only alter outwardly observable traits, but also affect gene expression profiles [10–13]. Natural selection can then act on this revealed variation either by selecting against dysregulated loci or, more rarely, facilitating adaptation [14]. In support of the first scenario, Neanderthal-derived variants with large effects on gene expression appear to be missing in modern humans [15]. Similarly, in a contemporary baboon hybrid zone, levels of introgression are systematically reduced near genes where ancestry predicts gene expression levels [16]. These examples suggest that gene regulatory divergence is common between closely related species, unmasked in hybrid populations, and helps shape the evolutionary outcomes of hybridization.

However, most studies of gene regulatory divergence in hybrids have been conducted in laboratory crosses rather than natural hybrid zones. While these studies have been foundational for understanding the overall genetic architecture of gene expression divergence, lab crosses differ from admixture in the wild in important ways. For example, laboratory crosses often focus on early generation hybrids of taxa that would not encounter each other in nature and would be unlikely to interbreed if they did [17–19]. These differences may account for more frequent evidence for gene regulatory dysregulation in the lab than in natural hybrids. Furthermore, studies in the lab do not clarify the evolutionary consequences of admixture for population variation in gene regulation in nature.

In this study, we set out to investigate the role of admixture and hybridization in shaping the genetic architecture of DNA methylation, a key gene regulatory mechanism, in a natural nonhuman primate hybrid zone. To do so, we paired genome-scale DNA methylation profiling with genome-wide resequencing data and local ancestry analysis for hundreds of individuals within a single admixed baboon population (yellow baboon (*Papio cynocephalus*) x anubis baboon (*P. anubis*)) [16,20–22]. We also map variants associated with DNA methylation variation in this population (i.e., methylation quantitative trait loci: meQTL) and investigate how they contribute to regulatory divergence between species and trait variance in the hybrid population.

We asked three sets of questions: (i) To what degree does genetic ancestry explain variation in DNA methylation patterns, and do ancestry effects in hybrid individuals recapitulate differences between unadmixed anubis and yellow individuals? (ii) Are changes in allele frequencies at meQTL (as opposed to distinct meQTL effect sizes) able to explain ancestry-associated DNA methylation? (iii) Does admixture lead to phenotypic variation visible to selection—and specifically, does it introduce new, functionally relevant regulatory variants? To address this last question, we use an experimental massively parallel reporter assay to test whether ancestry-associated CpG sites have the capacity to causally drive differences in enhancer activity. Together, our findings improve our understanding of the role of hybridization in shaping the genetic architecture of, and evolutionary potential for, gene regulation in natural populations.

## Results

### Local, but not global, ancestry is associated with variation in DNA methylation levels

Our study subjects were 256 individually recognized baboons living in the Amboseli ecosystem of Kenya, in an intensively monitored population that has been followed continuously since 1971 by the Amboseli Baboon Research Project (Table S1) [23]. This population falls in a natural hybrid zone in southern Kenya with predominantly yellow baboon ancestry, but where all individuals have some ancestry from anubis baboons (also referred to as olive baboons), a species that diverged from yellow baboons ∼1.4 million years ago [21,22,24]. Our primary data sets were genome-wide DNA methylation data profiles for 295 samples (38 animals were sampled multiple times) [25–27] and estimates of local genetic ancestry derived from whole-genome resequencing for all individuals [16] (Methods). We profiled DNA methylation at 2,218,202 autosomal CpG sites across the population, which followed expected genome-wide distributions and DNA methylation levels for RRBS data (Fig S1).

To concentrate on CpG sites with appreciable variation in genetic ancestry and DNA methylation levels, we focused on the 636,955 autosomal CpG sites that were not constitutively hyper- or hypomethylated in our sample (mean methylation ∈ [0.1,0.9]) and where each possible local ancestry state overlapping the CpG site (homozygous yellow, heterozygous, and homozygous anubis) was represented by at least 10 individuals. For each site, we modeled variation in DNA methylation levels as a function of local and global ancestry while controlling for technical and biological covariates known to affect DNA methylation (Methods). Consistent with previous findings for gene expression in this population and in admixed humans [16,28], we found little evidence for an association between genome-wide genetic ancestry and gene regulation (minimum FDR in this study = 58%), but substantial evidence for the effects of local genetic ancestry. Specifically, local genetic ancestry was significantly associated with DNA methylation levels at 100,482 CpG sites (FDR < 10%; 15.8% of tested sites; Table S2). Because some CpG sites present in the anubis baboon genome are absent in yellow baboons, local ancestry effects were asymmetric, with higher methylation in anubis alleles at 60% of detectable ancestry-associated CpG sites (see Supplementary Methods). We observed little to no evidence for non-additive effects (Fig S2; Methods), which is also in line with previous analyses of local ancestry and gene expression in this population [16]. Ancestry effects on DNA methylation in our sample therefore appear to be overwhelmingly additive and *cis*-acting.

### Ancestry-associated differential methylation in admixed Amboseli baboons recapitulates divergence in DNA methylation levels between species

Previous work investigated divergence in DNA methylation levels across the baboon radiation, including 9 anubis baboons (7 from Washington National Primate Research Center and 2 from Maasai Mara National Reserve, Kenya) and 6 yellow baboons from Mikumi National Park, Tanzania [29]. If DNA methylation differences between these populations reflect *cis*-regulatory genetic effects, as opposed to environmental or genetic background effects, they should be recapitulated in admixed animals within Amboseli. Focusing on 393,698 sites measured in both datasets, we detected a large number of CpG sites that were significantly differentially methylated between species, despite the modest sample size (Fig S3). 9,544 CpG sites were differentially methylated at a nominal p-value ≤ 0.01 (FDR ≈ 30%, where 3,937 sites would be expected by chance), including 480 sites that were significantly differentially methylated at an FDR of 10%.

Among the 9,544 sites with a nominal p-value ≤ 0.01, the estimated difference in DNA methylation between yellow and anubis baboons was correlated with the estimated difference between alleles within the Amboseli population (r = 0.695, *p* < 10^-300^). Consequently, if unadmixed anubis baboons exhibited higher DNA methylation levels than unadmixed yellow baboons in the between-species comparison, anubis alleles predicted higher DNA methylation levels relative to yellow alleles in Amboseli hybrids (and vice-versa; Fig 1B). CpG sites that differed between unadmixed yellow and anubis baboons were therefore also more likely to be associated with local ancestry in Amboseli — approximately 4-fold more than expected by chance (3,970 sites at a 10% FDR; Fisher’s Exact Test: log_2_(OR) = 2.00, *p* < 10^-300^). Among this set of sites (i.e., those that differed both between populations and by local ancestry within Amboseli), the estimated difference between anubis and yellow populations and the estimated difference between anubis and yellow alleles were highly correlated (r = 0.846, *p* < 10^-300^) and nearly universally (94.2%) directionally concordant. Evidence of overlapping effects was even stronger for the 480 CpG sites passing a 10% FDR in the between-species analysis. 330 of these sites (69%) were also significantly associated with local ancestry within Amboseli (log_2_(OR) = 4.36, *p* = 1.17x10^-148^). These results support a genetic basis for many interspecific differences in DNA methylation in baboons, robust enough to be detected despite major differences in geography, ecology, and sampling design between individuals sampled in distinct populations.

**Figure 1:**
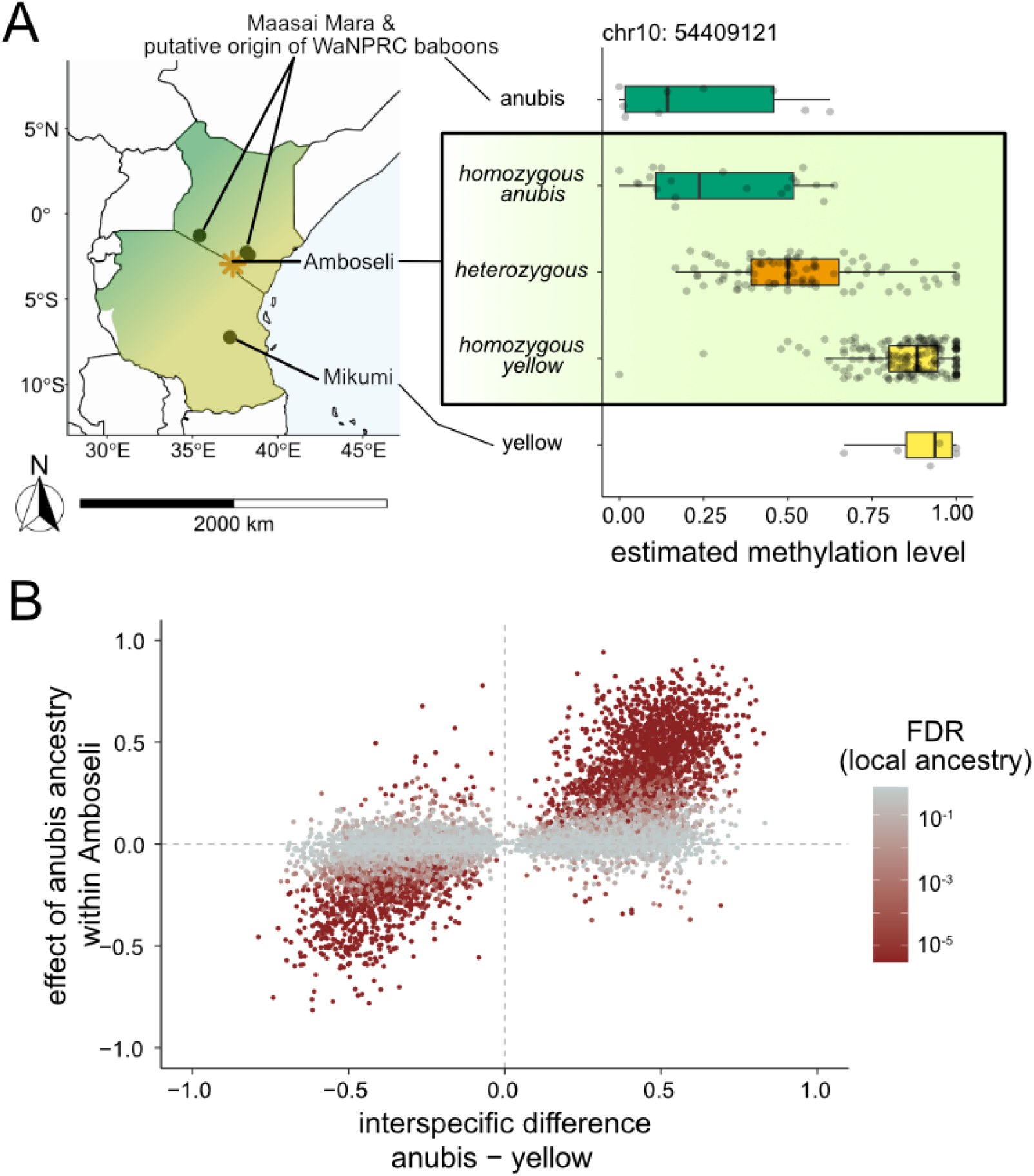
Associations between genetic ancestry and DNA methylation in a hybrid population converge with estimates of differential methylation between species. A. Geographic origins for baboons in this study (orange star = Amboseli), and DNA methylation levels at an example CpG site on chromosome 10 where variation in DNA methylation is both associated with local genetic ancestry and recapitulates the difference in DNA methylation between unadmixed yellow baboons and anubis baboons. B. Differences in DNA methylation associated with local ancestry in the Amboseli population are strongly correlated with differences in DNA methylation between unadmixed yellow and anubis baboons (r = 0.695, *p* < 10^-300^). Sites shown pass a nominal threshold of p < 0.01 for between population differences (n = 9,544 CpG sites). The x-axis represents the mean difference in DNA methylation between baboon species, where samples were collected from distinct populations; the y-axis represents the mean difference between yellow and anubis alleles segregating within Amboseli. Darker shades of red represent CpG sites with stronger evidence (smaller FDR) for ancestry-associated methylation within Amboseli.

### Allele frequency differences at methylation quantitative trait loci account for ancestry effects on DNA methylation

Anubis baboons and yellow baboons still share substantial genetic variation, although the frequencies of shared variants have diverged (F_ST_ estimates range between 0.23 and 0.33 [22,24]). Consequently, ancestry-associated differential methylation may reflect differentiation in the allele frequencies of genetic variants that influence DNA methylation levels (i.e., methylation quantitative trait loci, or closely linked sites) rather than changes in effect size at these sites or different causal sites entirely.

To test this hypothesis, we mapped meQTL in the Amboseli population for 172,044 SNP-CpG pairs (Methods). We identified 21,732 meQTL pairs (FDR < 10%). CpG sites disrupted by a SNP exhibited the expected reduction in DNA methylation levels, such that 96% of SNPs overlapping a CpG site were meQTL associated with reductions in DNA methylation levels (6,014 out of 6,264 sites). The minority of cases in which disrupted CpG sites were not associated with a meQTL (4% of pairs tested) were explained by low levels of DNA methylation even when the CpG site was intact (Supplementary Methods). Even excluding disrupted CpG sites, meQTL were more common when SNPs fell close to their associated CpG site (logistic regression β = -0.0092, *p* = 3.73x10^-132^), possibly because SNPs close to a focal CpG site are more likely to impact transcription factor binding [30]. 20,109 ancestry-associated CpG sites were tested as part of at least one CpG-SNP pair, 49% of which (9,843 CpG sites) were associated with a meQTL (FDR < 10%). The strong enrichment of meQTL at local ancestry-associated CpG sites (log_2_(OR) = 3.31, *p* < 10^-300^) suggests that divergence in meQTL frequencies may indeed underlie the association between genetic ancestry and DNA methylation.

We therefore asked whether meQTL and allele frequency differences between species could be combined to predict the association between genetic ancestry and DNA methylation. Here, we focused on the 9,843 CpG sites for which we detected both ancestry-associated DNA methylation in the Amboseli hybrid population (10% FDR) and an meQTL (10% FDR), and where allele frequency data were available at the meQTL SNP for yellow and anubis baboons outside the hybrid zone [16] (n= 9,136 unique SNPs). If *cis*-regulatory variants underlie the effects of local ancestry in the Amboseli population, the predicted local ancestry effect is expected to be the product of the estimated meQTL effect and the difference in allele frequencies between the parental taxa. Indeed, this simple calculation strongly predicts the observed local ancestry effect in the Amboseli sample (Spearman’s rho = 0.762, *p* = < 10^-300^; Fig 2). Thus, much of the DNA methylation divergence between yellow and anubis baboons can be explained by divergence in allele frequencies at large effect, *cis*-acting genetic variants associated with DNA methylation levels. Most of this divergence can be explained without requiring a contribution from epistasis or new functional variants, although our results do not exclude an additional contribution from these mechanisms.

**Figure 2:**
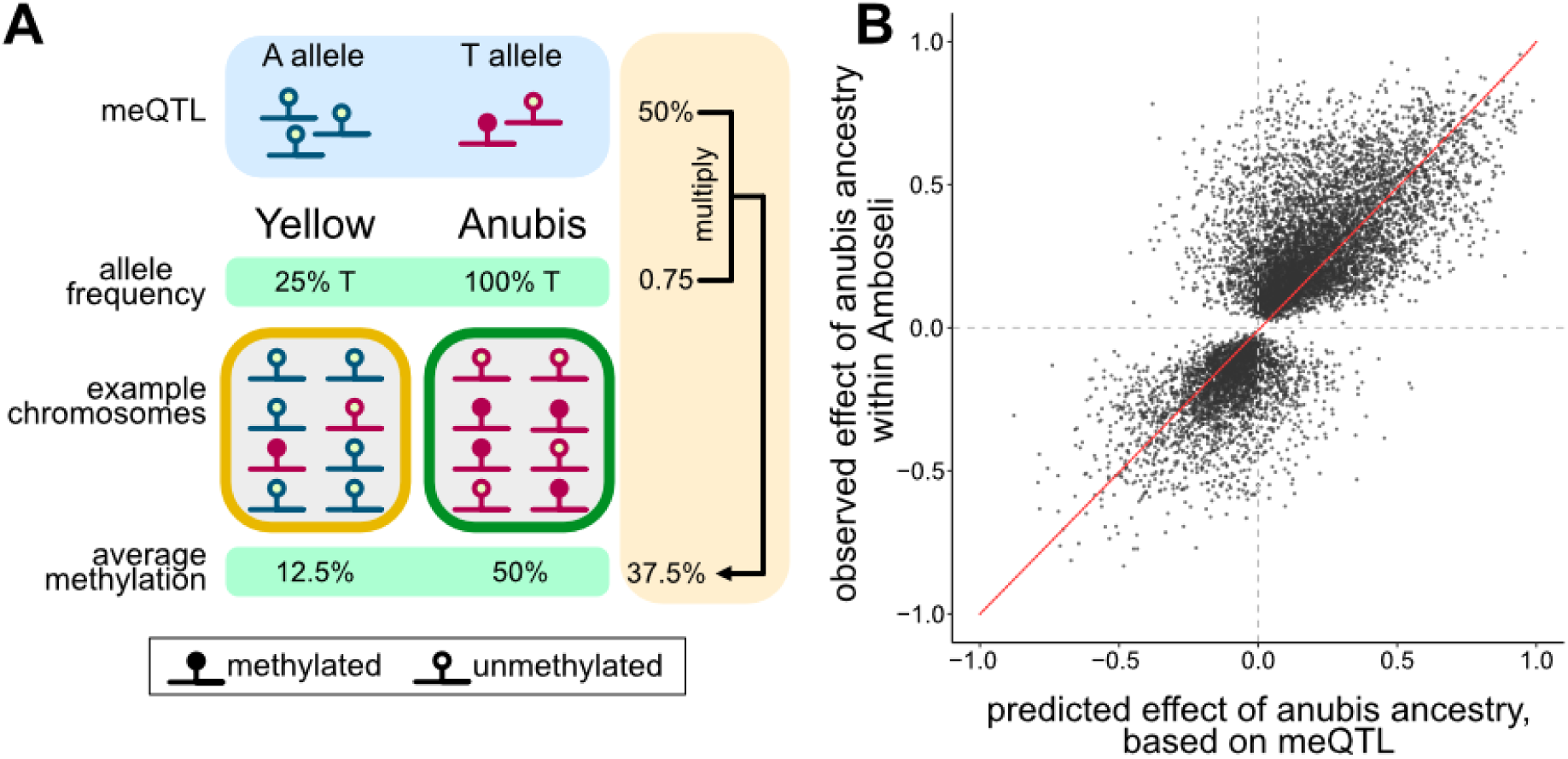
Ancestry-associated DNA methylation can be predicted from allele frequency differences at mapped meQTL. A. In the absence of epistasis or new functional variants in one or both lineages, ancestry differences in DNA methylation are predicted to be the product of allele frequency differences at a causal (or linked to causal) meQTL and that meQTL’s effect size. In this example, an meQTL is an A/T transversion and its effect is identical in yellow and anubis baboon populations. A alleles (blue) are always unmethylated while T alleles (red) are methylated 50% of the time (meQTL effect = 50%). T alleles are at 25% frequency in yellow baboons, producing a mean methylation level of 12.5% population-wide or within a pool of yellow alleles in a hybrid population. The T allele is fixed in anubis baboons, leading to a mean methylation level of 50% in anubis populations or within a pool of anubis alleles in a hybrid population. The ancestry effect (50% - 12.5% = 37.5%) is equivalent to the meQTL effect (50%) multiplied by the allele frequency difference (75%). B. For local ancestry-associated CpG sites, the estimated effect of anubis ancestry on DNA methylation levels in the Amboseli hybrids (y-axis) is predicted by the effect size of the meQTL for those sites and the difference in the meQTL allele frequency between yellow and anubis baboons (x-axis; Spearman’s rho = 0.762, *p* = < 10^-300^). Red line represents y=x.

### Admixture increases genetic variance for DNA methylation levels in the Amboseli baboon hybrid population

Our results above suggest that local ancestry effects on DNA methylation arise in part from changes in the allele frequencies of *cis*-acting meQTL. If that is the case, we predict that anubis admixture has increased both genetic and epigenetic variance in the Amboseli population. For meQTL SNPs, the minor allele frequency (MAF) increased 2.78 ± 15.4% (mean ± s.d.) when considering all individuals for whom resequencing data were available, compared to allele frequencies estimated only for locally homozygous yellow individuals (15,962 variants: paired t-test *p* = 9.48x10^-67^; Fig 3A). This effect is driven by meQTL that are at low frequency (≤10%) in yellow baboons but common in anubis baboons (6,518 variants: 12.5 ± 14.6% increase in MAF, *p* < 10^-300^; Fig S4). At the extreme, we identified 2,420 meQTL that were invariant among homozygous yellow individuals but variable in the population as a whole, representing regulatory variation that was introduced into the Amboseli population via admixture. These patterns are unlikely to be driven by kinship instead of ancestry, as allele frequency estimates are highly correlated between stretches of homozygous ancestry within Amboseli and unadmixed individuals sampled far from the hybrid zone (yellow ancestry vs yellow baboons: r = 0.911, *p* < 10^-300^; anubis ancestry vs anubis baboons: r = 0.781, *p* < 10^-300^).

**Figure 3:**
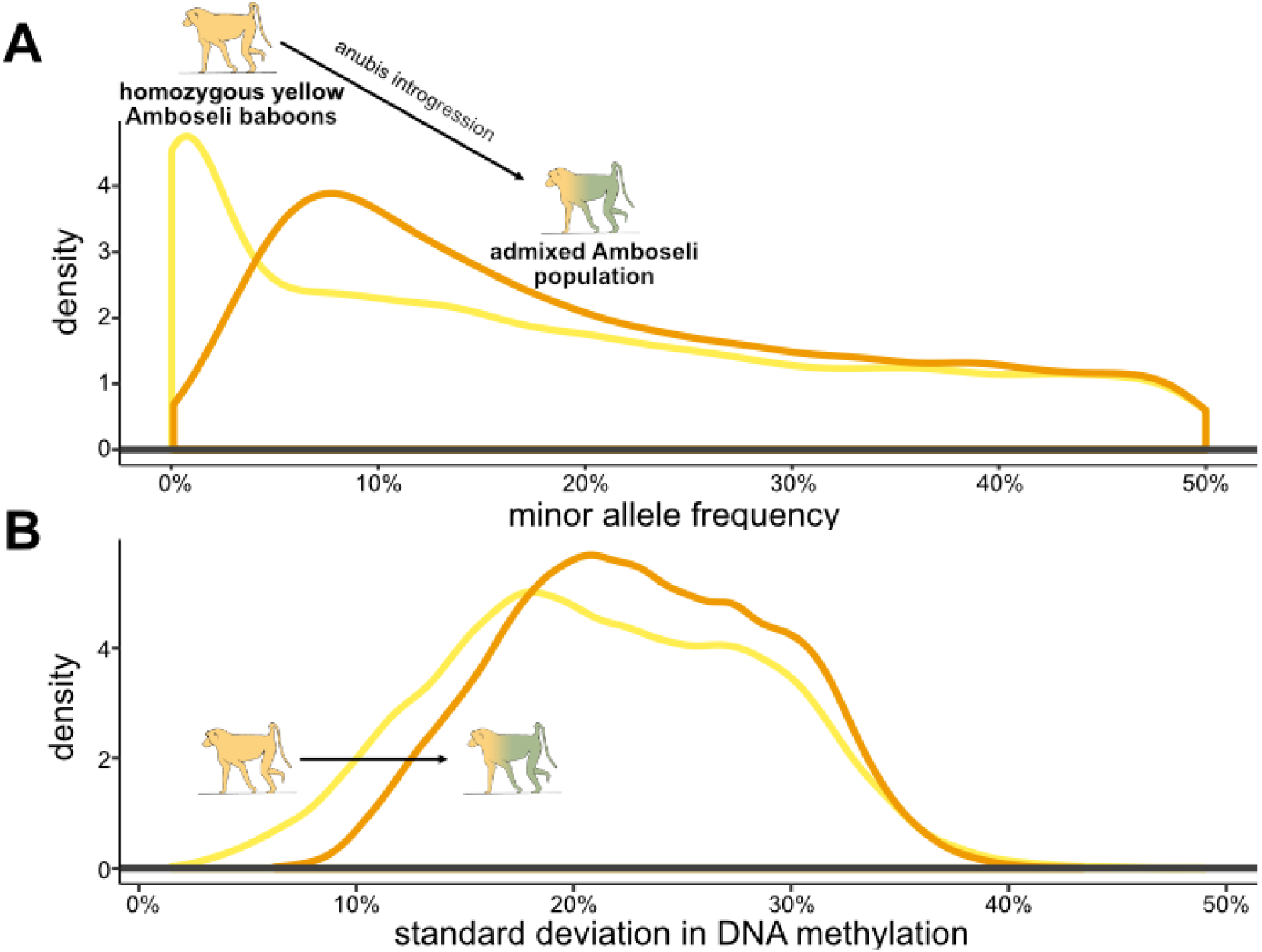
Admixture increases functional genomic variation within Amboseli. A. Density plot of the minor allele frequency at meQTL SNPs (n = 15,962 SNPs). Minor allele frequencies were calculated for all individuals with locally homozygous yellow baboon ancestry (yellow line) and for all genotyped individuals in the Amboseli population (orange line). There is a significant shift towards higher minor allele frequencies when including individuals with anubis ancestry (mean of 2.8%, paired t-test *p* = 9.48x10^-67^). Note that the minor allele frequency for the meQTL SNPs is shifted towards higher values than for all segregating variants, as there is greater power to detect meQTL at more common variants. B. Density plot of the standard deviation in DNA methylation levels, calculated from the ratio of methylated to total reads for each individual at each CpG site associated with an meQTL (n = 20,655). The curve shifts significantly to the right when calculated for all baboons in the Amboseli population (orange line) versus the subset with locally homozygous yellow baboon ancestry (yellow line) (mean increase of 19.7%, paired t-test p < 10^-300^).

These differences in meQTL allele frequencies translate into an increase in the phenotypic variance in DNA methylation in the admixed population. The average standard deviation of DNA methylation levels at a CpG site affected by a meQTL increases by 19.7% when comparing locally homozygous yellow individuals to the full population (n = 20,655 CpG sites, paired t-test p < 10^-300^; Fig 3B). This difference is magnified, as expected, for CpG sites that also exhibit a significant relationship between local ancestry and DNA methylation in Amboseli (n = 11,966 CpG sites): here, the standard deviation increases by 32.2% in the entire sample relative to locally homozygous yellow individuals (p < 10^-300^). In contrast, the increase is only 2.3% for CpG sites with an identifiable meQTL but no ancestry effect (n=8,689 CpG sites, p = 2.64x10^-4^). Within the set of sites with both significant local ancestry effects and mapped meQTL, the amount of added variance also increases with the amount of introgressed anubis ancestry at that locus, within the Amboseli population (linear model β = 0.28, *p =* 2.17x10^-5^).

### Functional consequences of local ancestry-associated CpG sites on gene regulation

Finally, one of the main motivations to study genetic effects on gene regulation in diverging species is to understand their downstream phenotypic consequences. For DNA methylation, this path would most likely proceed through changes in gene expression, but not all changes in DNA methylation are functionally important for shaping expression [31–33]. We therefore performed three analyses to investigate whether local ancestry-associated sites have the capacity to alter gene expression levels. First, we investigated the distribution of local ancestry-associated sites in the baboon genome. Ancestry-associated sites—especially the ones with the largest effect sizes—tended to be depleted in functionally important elements, including promoters, enhancers, and CpG islands (Fig 4A, with some differences between CpG sites disrupted by allelic variation versus those that are not: Fig S5). This observation is consistent with functional and sequence conservation in these elements. However, ancestry-associated CpG sites are modestly enriched in non-exonic parts of genes (i.e., untranslated regions and introns), suggesting possible effects on gene expression through regulatory elements contained in these regions of the genome.

**Figure 4.**
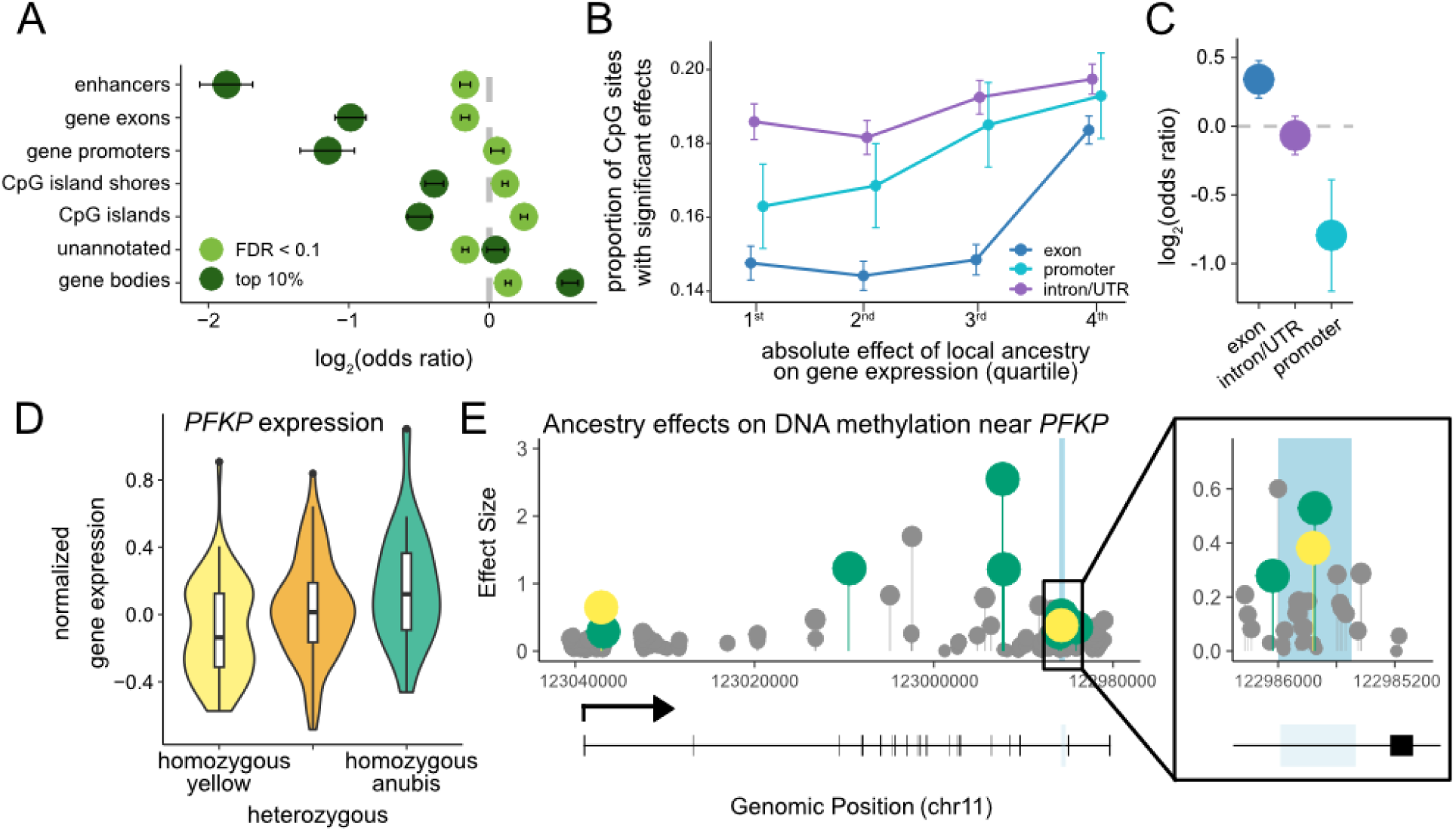
Functional consequences of ancestry-associated DNA methylation. A. Enrichment and depletion by genomic context for CpG sites in which local ancestry predicts DNA methylation levels, compared to the background set of all CpG sites analyzed. Results are shown for both all ancestry-associated CpG sites (n=100,482 sites, 10% FDR, light green) and for the most strongly associated sites (n=10,034, the top 10% of significant associations, dark green). Error bars represent the 95% confidence interval surrounding the log_2_(odds ratio). B. The proportion of CpG sites with ancestry-associated DNA methylation increases with the strength of the association between local ancestry and expression of the nearest gene (e.g., those in the most strongly associated fourth quartile; logistic regression *p* < 10^-6^). CpG sites are stratified by the functional context in which they occur (gene exons; promoters; or introns and untranslated regions (UTRs). Error bars represent the upper and lower 95% confidence intervals inferred from the standard error. Functional contexts are jittered to prevent overplotting. C. Enrichment and depletion of directionally concordant effect sizes between ancestry-associated CpG sites and nearby ancestry-associated genes. A positive log_2_(odds ratio) indicates that increased methylation is associated with increased expression. Error bars represent the 95% confidence interval surrounding the log_2_(odds ratio). D. Local ancestry effects on gene expression levels of *PFKP*. E. Effect of local ancestry on DNA methylation at CpG sites throughout the *PFKP* gene. Some effect sizes are greater than 1 because they are calculated following a logit transformation of the methylation values. Significant effects (FDR < 0.1) are colored based on the ancestry state that exhibits higher levels of DNA methylation (green = anubis ancestry, yellow = yellow baboon ancestry). CpG sites without significant associations are shown in gray. The light blue shaded area represents a region with methylation-dependent regulatory activity in the mSTARR-seq experiment. A gene diagram of *PFKP* is also included, with black boxes for exons and a bent arrow showing the transcription start site. A zoomed-in section of the PFKP gene (inset) shows the methylation-dependent enhancer experimentally identified in the mSTARR-seq assay, which also contains two ancestry-associated CpG sites. A third ancestry-associated site occurs just outside of this methylation dependent enhancer.

Second, we tested whether local ancestry-associated DNA methylation patterns are enriched near genes for which local ancestry has previously been correlated with gene expression levels in the Amboseli baboon population [16]. We considered 10,222 protein coding genes that were expressed in whole blood or white blood cells in RNA-seq data sets from this population [34–36]. DNA methylation was more likely to be significantly predicted by local ancestry when the CpG sites fell in the promoter (logistic regression β = 0.692, *p* = 3.27x10^-7^), exons (β = 0.788, *p* = 2.20x10^-111^), or untranslated regions (introns, 3’ UTR, 5’ UTR: β = 0.296, *p* = 8.09x10^-9^) of genes where local ancestry also predicted gene expression (Fig 4B). For example, a CpG site in the promoter region of a gene with strong ancestry-associated differential expression (top 10%) is 32% more likely to exhibit ancestry-associated DNA methylation levels than a CpG site in the promoter region of a randomly chosen expressed gene (20% more likely for a CpG site in an exon; 2.5% for a CpG site in an untranslated region). The direction of these effects was consistent with the canonical role of promoter DNA methylation as a silencing mark (Fig 4C). Specifically, for ancestry-associated CpG sites in the promoters of genes in the top quartile of ancestry effects on gene expression, increased DNA methylation was associated with decreased gene expression (log_2_(OR) = -0.794, *p* = 7.03x10^-5^). In contrast, increased DNA methylation predicted increased gene expression for ancestry-associated exonic CpG sites (log_2_(OR) = 0.342, *p* = 6.74x10^-7^) and there was no relationship between DNA methylation and gene expression for intronic or UTR ancestry-associated CpG sites (log_2_(OR) = 0.067, *p* = 0.341). These results indicate that local ancestry effects on DNA methylation and gene expression often—although not always—occur in tandem.

Finally, we integrated our local ancestry results with data from a massively parallel reporter assay, mSTARR-seq, to investigate whether sites detected in our scan for ancestry effects tend to causally alter enhancer-like activity, and hence gene expression output, when differentially methylated [31,37]. mSTARR-seq takes advantage of the ability to experimentally manipulate DNA methylation at CpG sites *in vitro*, on short DNA sequences that are identical other than their CpG methylation status (Fig S6). Cases in which the regulatory output from methylated sequences differ from unmethylated sequences point to functionally relevant, methylation-dependent regulatory activity. Cases in which methylated and unmethylated sequences do not differ, but drive higher regulatory activity than background, point to methylation-independent regulatory activity. Here, we drew on the results of an mSTARR-seq experiment applied to sequences from the baboon genome, which assayed 241,256 500-base pair windows [27] (∼4% of the genome, enriched for CpG-dense regions). Of these windows, 4,844 exhibited regulatory activity in the methylated state, unmethylated state, or both states (10% FDR). Enhancer activity was detected more frequently in the unmethylated condition (n = 4,335 regions detected at a 10% FDR versus n=1,717 in the methylated condition) and was sensitive to DNA methylation condition in 4,037 regions (10% FDR), most of which were more active in the unmethylated condition (87%, or 3,516 regions).

Among the 241,256 windows included in the mSTARR-seq experiment, 42,661 windows contained CpG sites also included in our analysis of local ancestry effects. In total, these windows contained 128,666 CpG sites and 20,218 (15.7%) of these sites exhibited associations between local ancestry and DNA methylation levels. Relative to all windows with tested CpG sites, cases of ancestry-associated DNA methylation in the Amboseli sample were depleted for evidence of regulatory potential (log_2_(OR) = -0.208, p = 0.0195). Conditional on falling in regions with enhancer-like activity in any state (methylated or unmethylated), they were also no more likely to exhibit methylation-dependence (log_2_(OR) = -0.087, p = 0.730). This result suggests that, overall, divergence in DNA methylation makes a limited causal contribution to gene expression divergence, even though ancestry-differentially methylated CpG sites and ancestry-differentially expressed genes often colocalize. This pattern agrees with generally higher levels of conservation in regulatory elements than background rates in the genome [38].

However, the combined data set also points to interesting candidates for further study, where DNA methylation is ancestry-associated in Amboseli and falls in regions where the mSTARR-seq data demonstrate causal effects of DNA methylation on gene expression. There are 258 such sites in our sample. These sites are the most likely of those we profiled to contribute to phenotypic differences between species and phenotypic diversity within the hybrid population. Of particular note are those that also fall near genes that are differentially expressed by local ancestry (n = 48 CpG sites near 14 ancestry-associated genes, Table S3). For example, anubis genetic ancestry is associated with higher expression of the gene *PFKP* (Fig 4D) and there are two CpG sites within a *PFKP* intron that also exhibit ancestry-associated DNA methylation (Fig 4E). *PFKP* encodes the rate-limiting step of glycolysis and genetic variation near *PFKP* is associated with body mass index and weight in humans [39]. *PFKP* expression also increases in captive baboons fed a high fat diet [40] and, in the Amboseli baboons, shows strong associations between DNA methylation and environmental variation in diet and resource base [26]. Our results here suggest that at least some variation in *PFKP* regulation also reflects genetic divergence between the two major extant baboon lineages. They therefore suggest an interesting candidate for studies of metabolic evolution in baboons, an important model system for human obesity [41,42].

## Discussion

Our results show how admixture influences the genetic architecture of DNA methylation in a natural primate hybrid zone. They emphasize the importance of genetic contributions to DNA methylation divergence, which in our system arise primarily through changes in the frequency of shared genetic variants in the parental taxa and result in largely additive, *cis*-acting local ancestry effects on DNA methylation levels. While such sites often occur near genes that exhibit ancestry-associated variation in gene expression, many sites that exhibit ancestry-associated DNA methylation may be functionally silent with respect to downstream gene regulation, at least in the cell types we interrogated here. However, based on our mSTARR-seq data, at least some ancestry-associated CpG sites are also functionally relevant for downstream gene regulation. At these loci, admixture increases the genetic contribution to DNA methylation variance, consistent with the idea that admixture affects hybrid zone dynamics by introducing new genetic and phenotypic variation [2]. Consequently, admixture in Amboseli increases the visibility of regulatory variation to natural selection.

Our findings thus provide new insight into the population and evolutionary genetic consequences of hybridization in primates. Unlike for new mutations, variation introduced through admixture often results in intermediate, instead of very rare, allele frequencies. Thus, introgressed variants can have immediate consequences for population-wide phenotypic variation and may be less likely to be lost through genetic drift [14,43]. Although introgressed variants are often selected against—potentially explaining why CpG sites with strong ancestry effects are depleted in mSTARR-seq-identified enhancers—beneficial introgressed alleles can rapidly increase in frequency [6,44]. To date, this phenomenon has been most clearly demonstrated for large-effect coding variants, such as those that affect coat color and wing pattern traits in dogs, snowshoe hares, and butterflies [45–47]. Our results suggest that one of the requirements for such scenarios—admixture-mediated expansion of trait diversity—is also commonly met for genetic variants underlying gene regulation. Data sets like the one produced here can therefore be intersected with sequence-based tests for selection to search for cases of adaptive introgression more broadly.

Consistent with previous studies of DNA methylation in naturally hybridizing animals [48–50], we observed almost entirely additive effects of local genetic ancestry. This observation differs from studies of gene regulation in laboratory crosses, which tend to identify widespread non-additivity [10–13,18]. Our findings are in line with the idea that reproductive isolation may be weaker in natural hybrid zones than in animal taxa chosen for laboratory experiments. An alternative possibility is that non-additive effects are more likely to arise for gene expression than DNA methylation. However, this explanation is inconsistent with other observations from our study system, where previous analyses of local ancestry effects on gene expression also identify mostly additive effects [16].

Previous studies have proposed a largely *cis*-regulatory basis for divergence in DNA methylation levels [50–51], including in natural hybrid zones (e.g., carrion crows and hooded crows in Europe [52]), which is consistent with our results. Indeed, we extend this argument by identifying *cis*-acting methylation QTL that, in combination with allele frequency differences between anubis and yellow baboons, are a remarkably good predictor of estimated local ancestry effects. 4,333 CpG sites where ancestry predicts DNA methylation also have a mappable, large-effect, putatively *cis*-acting meQTL in our data set. However, we were unable to identify meQTL for many ancestry-associated CpG sites, suggesting that many smaller effect meQTL remain to be discovered. Notably, our method for meQTL mapping, which measures DNA methylation and maps meQTL using allele-specific data from a single data set, is limited to the analysis of CpG-SNP pairs observed on the same read (in this data set, within 100 bp of each other). Improvements in population-scale genotyping, especially assisted by imputation [53], should substantially expand the scope of these types of analyses and alleviate several of the blind spots in studies like this one.

Nevertheless, our results already suggest that, although yellow and anubis baboons diverged more than a million years ago, meQTL effect sizes remain largely conserved across genetic backgrounds. This result extends findings in humans that differences in DNA methylation and gene expression between populations can be explained by QTL mapped within populations [54–56]. Indeed, recent studies indicate that even variants associated with complex traits such as height and gene expression often show conserved effect sizes across populations, suggesting that well-known challenges with the portability of genetic prediction stem primarily from differences in linkage disequilibrium and allele frequency [28,57–59]. Our findings suggest that, at least at the level of molecular phenotypes, effect sizes can remain consistent even over much longer timescales. This result raises interesting questions about the timescale of evolutionary divergence when effect sizes no longer tend to be shared. For example, to what degree does this timescale correlate with the timescale of reproductive isolation? And when genetic effect sizes are no longer shared, is this change due primarily to interactions with genetic background, interactions with fixed environmental differences, or simply the loss of shared genetic variation?

Finally, our findings have several practical implications for studies of gene regulatory evolution in natural populations. First, they indicate that genetic effect sizes may be partially generalizable across closely related species, which is important when one population or taxon is much more genetically well-sampled or well-characterized than others. The power and limits of this generalizability will be important to explore further. Second, our analyses support the value of comparative approaches for understanding gene regulatory evolution. Our study indicates that between-species differences from individuals sampled from environmentally and geographically distinct populations (and often in different sampling efforts) can nevertheless faithfully capture genetic effects that are recapitulated in the “common garden” of a naturally occurring hybrid zone. Finally, our results point to the feasibility of linking ancestry-associated DNA methylation and ancestry-associated gene expression within hybrid populations by combining observational mapping approaches with experimental assays to pin down causality. Future work should explore whether integrating sequence-based evidence for selection can further refine the discovery of variants responsible for reproductive isolation and/or trait divergence in non-model systems.

## Conclusion

This study shows that differences in allele frequencies at DNA methylation–associated variants account for a substantial portion of methylation divergence between closely related primate species. As a result, admixture increases genetic variation in DNA methylation in hybrid populations—and potentially, by extension, in other regulatory traits and their downstream effects. These findings provide new insight into how admixture shapes trait variation in natural hybrid zones. They also demonstrate the power of integrating comparative genomics, within-species QTL mapping, and functional assays to investigate gene regulatory evolution in non-model organisms.

## Methods

### Study population and samples

The primary subjects of this study were members of a long-term study population of wild baboons that has been monitored by the Amboseli Baboon Research Project (ABRP) for over five decades [23]. Approximately a third of the population directly descends from anubis or yellow-anubis admixed individuals who immigrated into the study area starting in the early 1980s. The remainder also exhibit lower levels of anubis admixture from older, unobserved bouts of gene flow [16,22].

For this study, we focused on 256 Amboseli individuals for whom both genome resequencing data and genome-wide DNA methylation data were available. The DNA methylation data were generated using reduced representation bisulfite sequencing (RRBS [60], see Supplementary Methods) and were previously analyzed in [25–27]. We also used genotype information for each of these individuals and an additional 178 individuals to calculate allele frequencies within the Amboseli population and to call local genetic ancestry, as in [16] (see Supplementary Methods).

### Ancestry effects on DNA methylation

RRBS data were processed as in [27] and mapped to the *Panubis1* anubis baboon genome [61] (see Supplementary Methods). The DNA methylation level for each individual at each CpG site was calculated as the proportion of reads with non-bisulfite converted (i.e., methylated) cytosine bases to total reads covering that site. We retained estimates for DNA methylation at 2,218,203 autosomal CpG sites. To focus on the sites most likely to exhibit biologically meaningful variation, we further excluded constitutively hypermethylated (mean DNA methylation level >0.90) and constitutively hypomethylated (mean DNA methylation level <0.10) sites. We also determined the genetic ancestry (local number of anubis alleles) for each individual at each CpG site (see Supplementary Methods) and retained CpG sites where our data set contained at least 10 individuals for each ancestry state (homozygous yellow, heterozygous, and homozygous anubis) at that CpG site (n = 636,955 analyzed CpG sites).

To test for associations between DNA methylation and genetic ancestry, we modeled the number of methylated reads as a function of the number of observed reads, local anubis ancestry, and global anubis ancestry while controlling for individual age, sex, dominance rank, early life adversity (measured using cumulative early life adversity and habitat quality, which are known to influence DNA methylation within this population [27]), bisulfite conversion rate, sequencing batch, and a random effect capturing genetic relatedness and population structure between samples. These models were run using MACAU (v4), which directly models observed count data and controls for population structure using a beta-binomial mixed effects model [62,63]. False discovery rates were estimated using a q-value approach calibrated against a permutation-based null model, implemented using the R *qvalue* package v2.24.0 [64,65]. Briefly, we compared the observed p-values to permutation-based empirical null distributions in which either local ancestry status (to test for local ancestry effects) or global ancestry (to test for global ancestry effects) was randomized across individuals while keeping all other information constant.

### Differences in DNA methylation between baboon species

To identify CpG sites where DNA methylation significantly differs between yellow and anubis baboons, we re-analyzed RRBS DNA methylation data on unadmixed yellow baboons from Mikumi National Park, Kenya (n = 6) and unadmixed anubis baboons from Maasai Mara National Reserve, Kenya (n = 2) and the Washington National Primate Research Center (n = 7) from [29]. We retained CpG sites covered in at least 5 individuals of each species, with mean coverage greater than 5x and with mean methylation (calculated across all individuals) between 0.1 and 0.9. We then tested each CpG site for a species difference in DNA methylation using MACAU [62] while controlling for the bisulfite conversion rate, which varied slightly between sequencing batches (Table S1). As above, we estimated the false discovery rate compared to an empirical null based on permuting the vector of species labels across samples.

### Genetic effects on DNA methylation

To identify genetic effects on DNA methylation, we jointly called DNA methylation levels and genotypes at nearby genetic variants from RRBS data for each sample (see Supplementary Methods). We retained positions where a genotype was called in at least 50% of individuals with an estimated minor allele frequency ≥ 0.05 in the present data set (to ensure power to map meQTL), and where the variant was known to be polymorphic within the Amboseli baboon population based on whole genome resequencing data [16].

We then extracted allele-specific counts for the number of methylated and unmethylated reads and excluded CpG sites with mean coverage less than 5x, mean methylation levels below 10%, or mean methylation levels above 90% (see Supplementary Methods). We also annotated disrupted CpG-SNP pairs where the SNP overlapped the CpG site.

Our dataset included 218,047 SNP-CpG pairs, spanning 74,967 unique SNPs and 169,812 unique CpG sites. For each SNP-CpG pair, we tested whether the SNP predicted methylation level using *IMAGE* (Fan, et al. 2019). *IMAGE* jointly models an additive effect of genotype on DNA methylation across individuals, along with the difference between alleles within heterozygotes (i.e., allele-specific DNA methylation). We calculated the false discovery rate via comparison to permuted data sets: first by randomizing genotypes among samples and then, within each heterozygote, randomly assigning each read to either the reference or alternate allele.

### Functional element annotations

We used gene annotations from NCBI to define coding sequences (https://ftp.ncbi.nlm.nih.gov/genomes/all/GCF/008/728/515/GCF_008728515.1_Panubis1.0/GCF_008728515.1_Panubis1.0_genomic.gtf.gz) and defined promoters as the 10 kb upstream of the 5’-most annotated transcription start site. Because experimentally derived baboon enhancer annotations are not available, we defined putative baboon enhancers by projecting coordinates from ENCODE H3K4me1 ChIP-seq data for human peripheral blood mononuclear cells [66] onto the *Panubis1.0* genome using the UCSC Genome Browser *liftover* tool [67] and a liftover file that we generated previously [16] (https://zenodo.org/records/5912279/files/Panubis1.0_to_hg38.chain.gz). CpG islands were annotated using the *emboss* cpgplot function with default parameters to identify windows of the genome longer than 200 bases with greater than 50% GC content and an observed/expected CpG ratio > 0.6 [68]. CpG island shores were defined as the 2 kb regions flanking either side of a CpG island [69]. Bed files for each annotation are available at https://doi.org/10.5281/zenodo.5912279. We then assigned CpG sites to one or more genomic contexts if they fell within the designated genomic regions. Finally, we tested for enrichment or depletion of differentially methylated sites using Fisher’s exact tests.

### CpG sites near genes with ancestry-associated expression

To test whether local ancestry-associated DNA methylation was more common near genes for which local ancestry is also associated with gene expression, we combined the analysis of DNA methylation presented here with ancestry effects on gene expression reported by [16]. Briefly, we drew on summary statistics from a prior analysis using RNA-seq gene expression data from 157 samples (n=10,202 genes), representing 145 unique individuals [34–36]. As previously reported, all samples were collected from reproductively mature individuals, and models for local ancestry effects controlled for age, sex, global genetic ancestry, genetic relatedness, and batch/sampling effects; for part of the data, collected on white blood cells, the analysis also controlled for cell type composition. Local ancestry was defined as a window spanning the gene transcription start site to transcription end site, plus an additional flanking 10 kb on each side. For very short genes, we extended the total window size to 50 kb, as local ancestry calls are less reliable for short windows. We used logistic regression to test whether the probability of detecting ancestry-associated DNA methylation is predicted by the absolute effect of local ancestry on gene expression.

### mSTARR-seq experiment and data analysis

Finally, to test whether local ancestry-associated DNA methylation can exert causal effects on enhancer activity, we used mSTARR-seq data previously analyzed in [27]. Briefly, plasmid libraries were prepared for mSTARR-seq transfection using genomic DNA isolated from the same anubis baboon used to construct the *Panubis1.0* genome assembly [61]. Genomic DNA was prepared using a Covaris S220 Focused-Ultrasonicator and via restriction enzyme digest with *Msp1* (in two separate batches that were then combined), followed by size selection for fragments between 300 and 800 bp. Size-selected fragments were cloned into *pmSTARRseq1* (the CpG-free plasmid backbone used for mSTARR-seq assays) using Gibson assembly. Methylated libraries were created via treatment of the resulting library with *M.Sss1*, which methylates all CpG sites on the inserted fragments. Unmethylated libraries underwent a parallel treatment with buffer only. Methylated and unmethylated plasmid libraries were transfected in parallel into human K562 cell lines, with six replicates per treatment. After 48 hours, cells were harvested and plasmid-specific DNA and RNA were extracted and sequenced following the protocol reported in [31].

For each replicate (n=6 unmethylated DNA, n=6 methylated DNA, n=6 unmethylated RNA, and n=6 methylated RNA), we used *bedtools2* [70] to count the number of reads that overlapped discrete, non-overlapping 500 bp windows of the baboon genome. For downstream analyses, we retained only windows with (i) median coverage greater than 4x in both methylated and unmethylated DNA samples, (ii) that were present in at least half of the DNA samples for both the methylated and unmethylated conditions (i.e., where there was sufficient DNA input to potentially drive gene expression), and (iii) where there were non-zero counts in at least half of RNA-seq replicates in either treatment (n=241,256 windows). Before testing for enhancer activity, library sizes were normalized with *calcNormFactors* from the *edgeR* package [71] and RNA-seq samples were normalized against DNA-seq samples using *voomWithQualityWeights* from the *limma* R package [72]. RNA abundance was modeled relative to DNA abundance, nested within treatment condition (methylated or unmethylated), as in [27,37]. Regions capable of regulatory activity generate more RNA than expected based on the DNA input. However, most regions exhibit no regulatory activity and therefore show many more input DNA reads than RNA reads. We therefore retained only regions with a positive effect for the sample type (RNA vs DNA) in either the methylated condition, unmethylated condition, or both.

To estimate the false discovery rate, we compared the observed results to empirical null distributions generated using 100 permutations of RNA/DNA labels within each pair of samples using the q-value approach [64,65]. While most observed windows have systematically less activity in the RNA condition than the DNA condition, permuted data have more balanced effect sizes. To recreate the criteria for positive effect sizes, for each permutation, we subsampled to retain the same number of regions with positive RNA vs DNA effect sizes as detected in the empirical sample. These p-values from these windows were used to generate the empirical null.

To identify methylation-dependent enhancer activity, we focused on the subset of windows (n=4,844) with regulatory activity in the methylated condition, unmethylated condition, or both. For these windows, we tested whether the effect of sample type (RNA vs DNA) differed between the methylated and unmethylated conditions (i.e. an interaction effect between RNA/DNA condition and the methylated/unmethylated condition). To estimate the false discovery rate for identifying methylation-dependent windows, we calculated q-values by comparing p-values from the empirical results against results from 100 permutations where treatment condition (methylated vs unmethylated) was randomly assigned to each DNA-RNA replicate pair.

## Declarations

### Ethics Approval

The Kenya-based research reported here was conducted with permission from the Kenya Wildlife Service (KWS), the National Environmental Management Authority (NEMA), and the National Commission for Science, Technology and Innovation (NACOSTI), which we currently renew on an annual basis. Samples were collected and exported under NEMA permits and CITES permits from KWS. We also hold a Memorandum of Agreement (MoA) with KWS, the University of Nairobi, the Kenya Institute of Primate Research, and the National Museums of Kenya that details the ABRP research mission and commitment to benefits sharing under the Nagoya Protocol. The research in this study was approved by the Institutional Animal Care and Use Committees at Duke University and the University of Notre Dame and the Ethics Council of the Max Planck Society and adhered to the laws and guidelines of the Kenyan government.

### Data and Materials Availability

The data supporting the conclusions of this article are available on the NCBI Sequence Read Archive. Previously generated DNA methylation data are available at PRJNA648767, SRP058411, and PRJNA970398. Previously published resequencing data are available under PRJNA755322, PRJNA308870, PRJNA433868, PRJNA54005, PRJNA20425, PRJNA162517, PRJNA54003, PRJNA54009, PRJNA54007, PRJNA251424, and PRJNA295782. Processed genotype data are available at https://zenodo.org/records/5912279. Previously published RNA-seq data are deposited under PRJNA269070, PRJNA480672, and PRJNA731520. Previously published mSTARR-seq data are available at PRJNA871297. Code for all results is available at https://github.com/TaurVil/Vilgalys_Amboseli_Methylation/.

## Author contributions

T.P.V. and J.T. conceived and designed the study. A.S.F., J.A.A., E.A.A., S.C.A., and J.T. collected data. T.P.V., A.S.F., and J.A.A. processed data. T.P.V. performed analyses. J.A.A. and D.L. conducted and contributed the mSTARR-seq experiments. T.P.V. and J.T. interpreted analyses. T.P.V. and J.T. wrote the manuscript with edits and revisions from all other coauthors.

## Acknowledgements

We thank J. Altmann for her fundamental contributions to research on the Amboseli baboons. We thank the University of Nairobi, Kenya Institute of Primate Research, the National Museums of Kenya, the Amboseli-Longido pastoralist communities, the Enduimet Wildlife Management Area, Ker & Downey Safaris, Air Kenya, and Safarilink for assistance in Kenya; R. Mututua, S. Sayialel, J.K. Warutere, I.L. Siodi, G. Marinka, B. Oyath, A. Meloimet, T. Wango, and V. Oudu as long-term members of the Amboseli Baboon Research Project (ABRP) team; K. Pinc, N. H. Learn, and J. B. Gordon for contributions to the ABRP database; and T.P. Voyles for managerial support in the wet lab.

## References

1. Barton NH, Hewitt GM. Analysis of Hybrid Zones. Annual Review of Ecology and Systematics. 1985;16:113–48.

2. Arnegard ME, McGee MD, Matthews B, Marchinko KB, Conte GL, Kabir S, et al. Genetics of ecological divergence during speciation. Nature. 2014;511:307–11.

3. Poelstra JW, Vijay N, Bossu CM, Lantz H, Ryll B, Müller I, et al. The genomic landscape underlying phenotypic integrity in the face of gene flow in crows. Science. 2014;344:1410–4.

4. Enbody ED, Sendell-Price AT, Sprehn CG, Rubin C-J, Visscher PM, Grant BR, et al. Community-wide genome sequencing reveals 30 years of Darwin’s finch evolution. Science. 2023;381:eadf6218.

5. Huerta-Sánchez E, Jin X, Asan, Bianba Z, Peter BM, Vinckenbosch N, et al. Altitude adaptation in Tibetans caused by introgression of Denisovan-like DNA. Nature. 2014;512:194–7.

6. Racimo F, Sankararaman S, Nielsen R, Huerta-Sánchez E. Evidence for archaic adaptive introgression in humans. Nat Rev Genet. 2015;16:359–71.

7. Quach H, Rotival M, Pothlichet J, Loh Y-HE, Dannemann M, Zidane N, et al. Genetic Adaptation and Neandertal Admixture Shaped the Immune System of Human Populations. Cell. 2016;167:643–656.e17.

8. Rinker DC, Simonti CN, McArthur E, Shaw D, Hodges E, Capra JA. Neanderthal introgression reintroduced functional ancestral alleles lost in Eurasian populations. Nat Ecol Evol. 2020;4:1332–41.

9. Wittkopp PJ, Kalay G. Cis-regulatory elements: molecular mechanisms and evolutionary processes underlying divergence. Nat Rev Genet. 2012;13:59–69.

10. Landry CR, Wittkopp PJ, Taubes CH, Ranz JM, Clark AG, Hartl DL. Compensatory *cis-trans* Evolution and the Dysregulation of Gene Expression in Interspecific Hybrids of Drosophila. Genetics. 2005;171:1813–22.

11. Wang L, Greaves IK, Groszmann M, Wu LM, Dennis ES, Peacock WJ. Hybrid mimics and hybrid vigor in *Arabidopsis*. Proc Natl Acad Sci USA. 2015;112. Available from: https://pnas.org/doi/full/10.1073/pnas.1514190112

12. Coolon JD, Wittkopp PJ. *cis-* and *trans* -Regulation in *Drosophila* Interspecific Hybrids. In: Chen ZJ, Birchler JA, editors. Polyploid and Hybrid Genomics. 1st ed. Wiley; 2013. p. 37–57.

13. Mack KL, Campbell P, Nachman MW. Gene regulation and speciation in house mice. Genome Res. 2016;26:451–61.

14. Barrett R, Schluter D. Adaptation from standing genetic variation. Trends in Ecology & Evolution. 2008;23:38–44.

15. McCoy RC, Wakefield J, Akey JM. Impacts of Neanderthal-Introgressed Sequences on the Landscape of Human Gene Expression. Cell. 2017;168:916–927.e12.

16. Vilgalys TP, Fogel AS, Anderson JA, Mututua RS, Warutere JK, Siodi IL, et al. Selection against admixture and gene regulatory divergence in a long-term primate field study. Science. 2022;377:635–41.

17. Runemark A, Moore EC, Larson EL. Hybridization and gene expression: Beyond differentially expressed genes. Molecular Ecology. 2024;e17303.

18. Behling AH, Winter DJ, Ganley ARD, Cox MP. Cross-kingdom transcriptomic trends in the evolution of hybrid gene expression. J of Evolutionary Biology. 2022;35:1126–37.

19. Frayer ME, Payseur BA. Do genetic loci that cause reproductive isolation in the lab inhibit gene flow in nature? Linnen C, Wolf J, editors. Evolution. 2024;78:1025–38.

20. Samuels A, Altmann J. Immigration of aPapio anubis male into a group ofPapio cynocephalus baboons and. Int J Primatol. 1986;7:131–8.

21. Alberts SC, Altmann J. Immigration and hybridization patterns of yellow and anubis baboons in and around Amboseli, Kenya. Am J Primatol. 2001;53:139–54.

22. Wall JD, Schlebusch SA, Alberts SC, Cox LA, Snyder-Mackler N, Nevonen KA, et al. Genomewide ancestry and divergence patterns from low-coverage sequencing data reveal a complex history of admixture in wild baboons. Molecular Ecology. 2016;25:3469–83.

23. Alberts SC, Altmann J. The Amboseli Baboon Research Project: 40 Years of Continuity and Change. In: Kappeler PM, Watts DP, editors. Long-Term Field Studies of Primates [Internet]. Berlin, Heidelberg: Springer Berlin Heidelberg; 2012. p. 261–87.

24. Rogers J, Raveendran M, Harris RA, Mailund T, Leppälä K, Athanasiadis G, et al. The comparative genomics and complex population history of Papio baboons. Science Advances. 2019;5:eaau6947.

25. Anderson JA, Johnston RA, Lea AJ, Campos FA, Voyles TN, Akinyi MY, et al. High social status males experience accelerated epigenetic aging in wild baboons. eLife. 2021;10:e66128.

26. Lea AJ, Altmann J, Alberts SC, Tung J. Resource base influences genome-wide DNA methylation levels in wild baboons (Papio cynocephalus). Molecular Ecology. 2016;25:1681–96.

27. Anderson JA, Lin D, Lea AJ, Johnston RA, Voyles T, Akinyi MY, et al. DNA methylation signatures of early-life adversity are exposure-dependent in wild baboons. Proc Natl Acad Sci USA. 2024;121:e2309469121.

28. Taylor DJ, Chhetri SB, Tassia MG, Biddanda A, Yan SM, Wojcik GL, et al. Sources of gene expression variation in a globally diverse human cohort. Nature. 2024;632:122–30.

29. Vilgalys TP, Rogers J, Jolly CJ, Baboon Genome Analysis, Mukherjee S, Tung J. Evolution of DNA Methylation in Papio Baboons. Mulligan C, editor. Molecular Biology and Evolution. 2019;36:527–40.

30. Shoemaker R, Deng J, Wang W, Zhang K. Allele-specific methylation is prevalent and is contributed by CpG-SNPs in the human genome. Genome Res. 2010;20:883–9.

31. Lea AJ, Vockley CM, Johnston RA, Del Carpio CA, Barreiro LB, Reddy TE, et al. Genome-wide quantification of the effects of DNA methylation on human gene regulation. Ponting CP, Weigel D, Aerts S, editors. eLife. 2018;7:e37513.

32. Maeder ML, Angstman JF, Richardson ME, Linder SJ, Cascio VM, Tsai SQ, et al. Targeted DNA demethylation and activation of endogenous genes using programmable TALE-TET1 fusion proteins. Nat Biotechnol. 2013;31:1137–42.

33. Kreibich E, Kleinendorst R, Barzaghi G, Kaspar S, Krebs AR. Single-molecule footprinting identifies context-dependent regulation of enhancers by DNA methylation. Molecular Cell. 2023;83:787–802.e9.

34. Anderson JA, Lea AJ, Voyles TN, Akinyi MY, Nyakundi R, Ochola L, et al. Distinct gene regulatory signatures of dominance rank and social bond strength in wild baboons. Philosophical Transactions of the Royal Society B: Biological Sciences. 2022;377:20200441.

35. Lea AJ, Akinyi MY, Nyakundi R, Mareri P, Nyundo F, Kariuki T, et al. Dominance rank-associated gene expression is widespread, sex-specific, and a precursor to high social status in wild male baboons. Proceedings of the National Academy of Sciences. 2018;115:E12163–71.

36. Tung J, Zhou X, Alberts SC, Stephens M, Gilad Y. The genetic architecture of gene expression levels in wild baboons. Dermitzakis ET, editor. eLife. 2015;4:e04729.

37. Johnston RA, Aracena KA, Barreiro LB, Lea AJ, Tung J. DNA methylation-environment interactions in the human genome. eLife. 2024;12:RP89371.

38. Andrews G, Fan K, Pratt HE, Phalke N, Zoonomia Consortium§, Karlsson EK, et al. Mammalian evolution of human cis-regulatory elements and transcription factor binding sites. Science. 2023;380:eabn7930.

39. Scuteri A, Sanna S, Chen W-M, Uda M, Albai G, Strait J, et al. Genome-Wide Association Scan Shows Genetic Variants in the FTO Gene Are Associated with Obesity-Related Traits. Barsh G, editor. PLoS Genet. 2007;3:e115.

40. Lin W, Wall JD, Li G, Newman D, Yang Y, Abney M, et al. Genetic regulatory effects in response to a high-cholesterol, high-fat diet in baboons. Cell Genomics. 2024;4:100509.

41. Cox LA, Comuzzie AG, Havill LM, Karere GM, Spradling KD, Mahaney MC, et al. Baboons as a Model to Study Genetics and Epigenetics of Human Disease. ILAR Journal. 2013;54:106–21.

42. VandeBerg JL, Williams-Blangero S, Tardif SD. The baboon in biomedical research. New York: Springer; 2009.

43. Slatkin M, Lande R. Segregation variance after hybridization of isolated populations. Genet Res. 1994;64:51–6.

44. Hedrick PW. Adaptive introgression in animals: examples and comparison to new mutation and standing variation as sources of adaptive variation. Mol Ecol. 2013;22:4606–18.

45. Cubaynes S, Brandell EE, Stahler DR, Smith DW, Almberg ES, Schindler S, et al. Disease outbreaks select for mate choice and coat color in wolves. Science. 2022;378:300–3.

46. Jones MR, Mills LS, Alves PC, Callahan CM, Alves JM, Lafferty DJR, et al. Adaptive introgression underlies polymorphic seasonal camouflage in snowshoe hares. Science. 2018;360:1355–8.

47. The Heliconius Genome Consortium. Butterfly genome reveals promiscuous exchange of mimicry adaptations among species. Nature. 2012;487:94–8.

48. Boman J, Qvarnström A, Mugal CF. Regulatory and evolutionary impact of DNA methylation in two songbird species and their naturally occurring F1 hybrids. BMC Biol. 2024;22:124.

49. von Holdt B, Heppenheimer E, Petrenko V, Croonquist P, Rutledge LY. Ancestry-Specific Methylation Patterns in Admixed Offspring from an Experimental Coyote and Gray Wolf Cross. Journal of Heredity. 2017;108:341–8.

50. Wang X, Werren JH, Clark AG. Allele-Specific Transcriptome and Methylome Analysis Reveals Stable Inheritance and Cis-Regulation of DNA Methylation in Nasonia. Barton NH, editor. PLoS Biol. 2016;14:e1002500.

51. Orozco LD, Rubbi L, Martin LJ, Fang F, Hormozdiari F, Che N, et al. Intergenerational genomic DNA methylation patterns in mouse hybrid strains. Genome Biol. 2014;15:R68.

52. Merondun J, Wolf JB. DNA methylation reflects cis-genetic differentiation across the European crow hybrid zone. Molecular Ecology. 2025: e70026.

53. Watowich MM, Chiou KL, Graves B, Montague MJ, Brent LJN, Higham JP, et al. Best practices for genotype imputation from low-coverage sequencing data in natural populations. Molecular Ecology Resources. 2023;1755–0998.13854.

54. Banovich NE, Lan X, McVicker G, Geijn B van de, Degner JF, Blischak JD, et al. Methylation QTLs Are Associated with Coordinated Changes in Transcription Factor Binding, Histone Modifications, and Gene Expression Levels. PLOS Genetics. 2014;10:e1004663.

55. Heyn H, Moran S, Hernando-Herraez I, Sayols S, Gomez A, Sandoval J, et al. DNA methylation contributes to natural human variation. Genome Res. 2013;23:1363–72.

56. Natri HM, Hudjashov G, Jacobs G, Kusuma P, Saag L, Darusallam CC, et al. Genetic architecture of gene regulation in Indonesian populations identifies QTLs associated with global and local ancestries. The American Journal of Human Genetics. 2022;109:50–65.

57. Cavazos TB, Witte JS. Inclusion of variants discovered from diverse populations improves polygenic risk score transferability. Human Genetics and Genomics Advances. 2021;2:100017.

58. Ding Y, Hou K, Xu Z, Pimplaskar A, Petter E, Boulier K, et al. Polygenic scoring accuracy varies across the genetic ancestry continuum. Nature. 2023;618:774–81.

59. Saitou M, Dahl A, Wang Q, Liu X. Allele frequency impacts the cross-ancestry portability of gene expression prediction in lymphoblastoid cell lines. The American Journal of Human Genetics. 2024;S0002929724003781.

60. Boyle P, Clement K, Gu H, Smith ZD, Ziller M, Fostel JL, et al. Gel-free multiplexed reduced representation bisulfite sequencing for large-scale DNA methylation profiling. Genome Biology. 2012;13:R92.

61. Batra SS, Levy-Sakin M, Robinson J, Guillory J, Durinck S, Vilgalys TP, et al. Accurate assembly of the olive baboon (*Papio anubis*) genome using long-read and Hi-C data. GigaScience. 2020;9:giaa134.

62. Lea AJ, Tung J, Zhou X. A Flexible, Efficient Binomial Mixed Model for Identifying Differential DNA Methylation in Bisulfite Sequencing Data. PLOS Genetics. 2015;11:e1005650.

63. Sun S, Hood M, Scott L, Peng Q, Mukherjee S, Tung J, et al. Differential expression analysis for RNAseq using Poisson mixed models. Nucleic Acids Research. 2017;45:e106–e106.

64. Storey JD, Bass AJ, Dabney A, Robinson D. qvalue: Q-value estimation for false discovery rate control. R package. 2021. Available from: http://github.com/jdstorey/qvalue

65. Storey JD, Tibshirani R. Statistical significance for genomewide studies. PNAS. 2003;100:9440–5.

66. The ENCODE Project Consortium. An integrated encyclopedia of DNA elements in the human genome. Nature. 2012;489:57–74.

67. Nassar LR, Barber GP, Benet-Pagès A, Casper J, Clawson H, Diekhans M, et al. The UCSC Genome Browser database: 2023 update. Nucleic Acids Research. 2023;51:D1188–95.

68. Larsen F, Gundersen G, Lopez R, Prydz H. CpG islands as gene markers in the human genome. Genomics. 1992;13:1095–107.

69. Irizarry RA, Ladd-Acosta C, Wen B, Wu Z, Montano C, Onyango P, et al. The human colon cancer methylome shows similar hypo- and hypermethylation at conserved tissue-specific CpG island shores. Nat Genet. 2009;41:178–86.

70. Quinlan AR, Hall IM. BEDTools: a flexible suite of utilities for comparing genomic features. Bioinformatics. 2010;26:841–2.

71. Robinson MD, McCarthy DJ, Smyth GK. edgeR : a Bioconductor package for differential expression analysis of digital gene expression data. Bioinformatics. 2010;26:139–40.

72. Ritchie ME, Phipson B, Wu D, Hu Y, Law CW, Shi W, et al. limma powers differential expression analyses for RNA-sequencing and microarray studies. Nucleic Acids Research. 2015;43:e47–e47.

