## Supplementary Information for "Admixture influences the genetic architecture of DNA methylation in a wild primate hybrid zone"

### Supplementary Methods

#### Study population and samples

Genomic data from baboons in the Amboseli study population were derived from blood samples collected by the Amboseli Baboon Research Project (ABRP) following well-established, minimally invasive procedures (Altmann et al., 1996; Lea et al., 2016; Tung et al., 2009, 2015). Briefly, animals were immobilized with a Telazol-loaded dart delivered through a hand-held blow gun and transferred to a processing site for blood sample collection. Following sample collection, study subjects were allowed to regain consciousness in a covered holding cage and then released near their social group and closely monitored. Blood samples were stored at the field site or at an ABRP-affiliated laboratory at the University of Nairobi until they were transported to the United States.

This study focuses on 256 Amboseli individuals for whom both genome resequencing data and genome-wide DNA methylation data were available. The DNA methylation data were generated using reduced representation bisulfite sequencing (RRBS: Boyle et al., 2012) and were previously analyzed in (Anderson et al., 2021, 2024; Lea et al., 2016). Genome resequencing data and local ancestry calls were available for all 256 individuals, as well as an additional 186 animals who we used to more precisely estimate allele frequencies in the Amboseli population. These data were first reported in Vilgalys et al., 2022. Finally, to estimate between-species differences in DNA methylation, we also drew on reduced representation bisulfite sequencing data for 6 yellow and 9 anubis baboons, first reported in Vilgalys et al., 2019.

#### RRBS data generation and processing

The primary Amboseli data set involved 256 unique baboons, several of whom were sampled multiple times, for a total of 295 RRBS libraries (Anderson et al., 2021, 2024; Lea et al., 2016) (**Table S1**). Each RRBS library was prepared from ~180 ng of whole blood-extracted genomic DNA. Libraries were barcoded and pooled together in sets of 10-12 samples, subjected to sodium bisulfite conversion using the EpiTect Bisulfite Conversion kit (QIAGEN), and then PCR amplified for 16 cycles prior to sequencing on an Illumina HiSeq platform. To assess the efficiency of the bisulfite conversion, 1 ng of unmethylated lambda phage DNA (Sigma Aldrich) was added to each sample prior to library construction. All sequences were trimmed for adapter contamination, RRBS end repair, and base quality using Trim Galore! (Babraham Bioinformatics).

For each RRBS sample, we mapped reads to the *Panubis1* anubis baboon genome (Batra et al., 2020) using BSMAP (Xi & Li, 2009). We used BSMAP's three-nucleotide mapping option, which eliminates most heterospecific mapping biases within *Papio* baboons (Vilgalys et al.,

2019). The DNA methylation level for each individual at each CpG site was calculated as the proportion of reads with unconverted (i.e., methylated) cytosine bases to total reads covering that site. Based on reads mapped to the lambda phage genome, all samples had a bisulfite conversion efficiency greater than 98% (**Table S1**).

We filtered for CpG sites where methylation levels were estimated for at least 100 samples and the CpG site had a mean coverage ≥ 5x (n = 2,218,202 autosomal CpG sites). DNA methylation levels across the genome followed expected patterns, including hypomethylation in CpG islands and promoters and hypermethylation in regions of the genome outside of genes and putatively functional regulaory elements. As expected for RRBS data, profiled sites were also strongly enriched near genes and CpG islands (**Figure S1**). At least one CpG sites was included in our final data set for 64.7% of promoters and 79.5% of protein-coding gene bodies in the *Panubis1.0* NCBI genome annotation (GCF_008728515.1_Panubis1.0).

### Local ancestry calling

We used local ancestry calls reported in Vilgalys et al., 2022. Briefly, local ancestry was called for each Amboseli individual relative to a reference panel of putatively unadmixed, wild-born yellow and anubis baboons used to found the Southwest National Primate Research Center baboon colony (n=24 anubis; n=9 yellow) (Robinson et al., 2019). Whole genome resequencing reads for each sample were mapped to the *Panubis1* genome (Batra et al., 2020) using *bowtie2* (Langmead & Salzberg, 2012). We then removed putative PCR duplicates using MarkDuplicates in *Picard* (Broad Institute 2019) and called genotypes using the Genome Analysis Toolkit (*GATK*) (McKenna et al., 2010). We retained biallelic SNPs that passed the following hard filters: maf > 0.05; QD < 2.0; MQ < 35.0; FS > 60.0; MQRank-Sum < -12.5; and ReadPosRankSum < -8.0. The same process was repeated for the reference panel individuals, for whom heterospecific ancestry was masked on a per-individual basis (see Vilgalys et al., 2022).

We estimated local ancestry for each Amboseli individual using *LCLAE* (Low Coverage Local Ancestry Estimation: Wall et al., 2016). This method estimates the number of anubis alleles at each SNP based on the genotype likelihoods for nearby sites (for autosomes only). Calls are therefore based on integrated information across spatially clustered variants, resulting in reliable assignments of ancestry from low coverage data (Vilgalys et al., 2022; Wall et al., 2016). To run *LCLAE*, we filtered for ancestry informative SNPs (defined as those with a 20% difference in allele frequency between the reference panel yellow and anubis baboons) that were called in both the reference panel and our set of Amboseli samples. We then calculated the most likely ancestry for each SNP based on genotype calls in the surrounding 35 kb window centered on the focal SNP and used majority rule across windows that included that SNP to assign a final ancestry call (i.e., 0, 1, or 2 anubis alleles). Finally, we collapsed contiguous SNPs that were assigned to the same ancestry state into ancestry tracts. We placed tract break points, where ancestry states switched, at the base pair exactly intermediate between SNPs assigned to different ancestry states. Based on evidence that short ancestry tracts are the most prone to error (Vilgalys et al., 2022), we removed ancestry tracts shorter than 1 kb.

### Non-additive ancestry effects on DNA methylation

We used piece-wise regression to test for non-additive effects of local ancestry on DNA methylation. To our knowledge, there is no available software that implements piecewise regression with count data and a mixed effects model. We therefore used DNA methylation level estimates rather than raw count data for this analysis. Specifically, we calculated the methylation ratio as the proportion of reads with methylated cytosine bases to total reads at each individual-CpG site combination and quantile-quantile normalized the resulting matrix of methylation ratios. While typically less powerful than directly modeling raw count data, approaches that analyze quantile-quantile normalized methylation ratios have similar power as count-based methods for relatively large sample sizes, such as that used here (Lea et al., 2015).

We then used ComBat from the *sva* R package (Leek et al., 2012) to remove batch effects between sequencing batches. To run ComBat, the dataset was split by chromosome and sequencing batch, and we imputed missing data for each chromosome-batch combination using the *impute* package in R (Hastie, 2017). We then used ComBat to remove the technical effects of sequencing batch, controlling for sample age, sex, and global anubis ancestry. Finally, we reintroduced NAs for site-individual combinations with no count data. We also regressed out the random effect estimated based on genetic relatedness from the batch-corrected data set using *EMMREML*. Finally, we fit piece-wise regression models for the effect of local ancestry on each CpG site, with a break point at ancestry state = 1 (heterozygous samples) using the *segmented* package in R (Muggeo, 2016). As for other analyses, we calculated false discovery rates via comparison against a permutation-based null distribution.

### Genetic effects on DNA methylation

To identify genetic effects on DNA methylation, we jointly called DNA methylation levels and genotypes at nearby genetic variants for each sample. Trimmed RRBS reads were mapped to the *Panubis1* genome using *bismark* (Krueger & Andrews, 2011). We then called SNPs for each individual using *CGmaptools*, a SNP calling program specifically designed for bisulfite sequencing data (Guo et al., 2018). We used modified source code that also outputs homozygous reference genotypes (in addition to heterozygous and homozygous alternate genotypes; Fan, et al. 2019) and filtered out SNPs that were ambiguous or called using fewer than three reads. After SNP calling, we merged variant calls and then retained only biallelic positions called in at least 50% individuals (including homozygous reference calls) with an estimated minor allele frequency ≥ 0.05.

We then used *CGmaptools* to obtain CpG-SNP pairs where the SNP and CpG site were profiled on the same sequencing read. We extracted the methylation level estimates for each CpG site in the form of the number of methylated read counts and the number of total read counts, at the individual level for homozygotes and for each allele separately for heterozygotes. Following (Fan et al., 2019), we excluded CpG sites measured in fewer than 20 individuals, with mean coverage less than 5x, or that were hyper- or hypomethylated in the population as a whole (retaining CpG sites with mean methylation ∈ [0.1,0.9]). We also annotated CpG sites that were disrupted by a nearby SNP (e.g., a SNP in the Amboseli population that overlapped the C or G position, which would abolish the CpG site).

This procedure resulted in a data set of 172,044 SNP-CpG pairs, spanning 57,406 unique SNPs and 140,301 unique CpG sites, which we then used to map meQTL using the software *IMAGE* (Fan, et al. 2019). *IMAGE* jointly models an additive effect of genotype on DNA methylation across individuals, along with the difference between alleles within heterozygotes (i.e., allele-specific DNA methylation). We calculated the false discovery rate via comparison to permuted data sets: first by randomizing genotypes among samples and then, within each heterozygote, randomly assigning each read to either the reference or alternate allele.

We identified 6,264 SNP-CpG pairs where the SNP overlaps the CpG site and abolishes the CpG motif (e.g., where a C>T mutation changes the CpG site to become TpG). Because alleles that no longer carry a C cannot possibly be methylated, heterozygotes for CpG/NpG should exhibit allele-specific methylation, as long as the original C allele is itself appreciably methylated. This is the pattern we observed: 96% of disrupted CpG sites are associated with meQTL (10% FDR), which increases to 98.4% when considering only CpG sites with greater than 20% mean methylation. In all cases, the allele containing the disrupted CpG site has decreased methylation.

### Increased DNA methylation associated with anubis alleles

As noted in the main text, we observed that anubis alleles tended to have higher methylation levels than the yellow allele for approximately 60% of ancestry-associated CpG sites. This phenomenon occurred when analyzing ancestry effects within Amboseli and when comparing anubis individuals to yellow individuals in our between-species comparison. Such an asymmetry is unlikely to be the result of mapping bias: when using three nucleotide mapping, anubis and yellow alleles are equally likely to map to the anubis reference genome, regardless of methylation status (Vilgalys et al. 2019). Instead, it likely emerges from cases in which CpG sites occur in the anubis reference genome—and are therefore included in the data set as estimable sites—but are rare or absent in yellow baboons, resulting in low or zero mean DNA methylation levels at those loci for individuals with locally yellow ancestry. Anubis alleles therefore appear more highly methylated because reference CpG sites are more likely to be present on anubis baboon haplotypes than yellow haplotypes.

To test this hypothesis, we focused on CpG sites (i) that exhibited ancestry-associated DNA methylation in Amboseli, and (ii) where segregating variation in Amboseli meant that the CpG site was sometimes disrupted (n = 41,219 CpG sites). The proportion of sites with higher methylation in anubis alleles increases as the disrupting allele becomes more common in yellow baboons than anubis baboons (logistic regression $p < 10^{-16}$). For example, at CpG sites where the disrupting allele is more than 50% more common in yellow baboons than anubis baboons, nearly all (99%) of CpG sites show higher DNA methylation at anubis alleles in the Amboseli hybrid population. This pattern is indeed symmetrical: at CpG sites where the disrupting allele is 50% more common in anubis baboons than yellow baboons, nearly all (98%) of CpG sites show higher DNA methylation at *yellow* alleles in the Amboseli hybrid zone. However, as a consequence of use of the anubis reference genome, our data set contains 10x as many sites where the CpG-disrupting variant is more common in yellow baboons than sites where the variant is more common in anubis baboons (7,204 versus 739). Thus, the observed bias in effect sizes emerges because we analyzed CpG sites that were intact in the reference genome (i.e., more common, where disrupting variants occur, in anubis haplotypes than yellow haplotypes).

Importantly, these instances represent *bona fide* cases of differential methylation. We are simply blind to their counterpart: CpG sites found more commonly on yellow haplotypes than anubis haplotypes, which would result in higher DNA methylation levels in yellow baboons if we had used a yellow baboon reference genome.

#### Admixture increases genetic variance for DNA methylation levels

When analyzing the relationship between admixture and genetic variance in DNA methylation levels, we considered the alternative possibility that any increase in the full, admixed data set, relative to homozygous yellow genotypes, could be produced by kinship instead of ancestry. Under this hypothesis, individuals with homozygous yellow ancestry are likely to be, on average, more closely related to each other than they are to individuals with anubis ancestry. If so, the differences we observed may be a product of sampling more diverse family groups rather than ancestry-associated changes in allele frequency. However, allele frequencies from an independent panel of unadmixed yellow baboons (Robinson et al., 2019; Vilgalys et al., 2022) were highly correlated with those estimated from homozygous yellow ancestry tracts within Amboseli (55,773 SNPs: $r = 0.895$, $p < 10^{-300}$). Furthermore, allele frequencies in Amboseli can be predicted as a function of the allele frequencies in unadmixed yellow and unadmixed anubis baboons and per-locus levels of anubis ancestry ($r = 0.88$, $p < 10^{-300}$). Thus, anubis introgression has increased both overall genetic diversity in Amboseli, including genetic diversity at variants associated with DNA methylation.

### References


Altmann, J., Alberts, S. C., Haines, S. A., Dubach, J., Muruthi, P., Coote, T., Geffen, E., Cheesman, D. J., Mututua, R. S., Saiyalel, S. N., Wayne, R. K., Lacy, R. C., & Bruford, M. W. (1996). Behavior predicts genes structure in a wild primate group. *Proceedings of the National Academy of Sciences*, *93*(12), 5797–5801. https://doi.org/10.1073/pnas.93.12.5797

Anderson, J. A., Johnston, R. A., Lea, A. J., Campos, F. A., Voyles, T. N., Akinyi, M. Y., Alberts, S. C., Archie, E. A., & Tung, J. (2021). High social status males experience accelerated epigenetic aging in wild baboons. *eLife*, *10*, e66128. https://doi.org/10.7554/eLife.66128

Anderson, J. A., Lin, D., Lea, A. J., Johnston, R. A., Voyles, T., Akinyi, M. Y., Archie, E. A., Alberts, S. C., & Tung, J. (2024). DNA methylation signatures of early-life adversity are exposure-dependent in wild baboons. *Proceedings of the National Academy of Sciences*, *121*(11), e2309469121. https://doi.org/10.1073/pnas.2309469121

Batra, S. S., Levy-Sakin, M., Robinson, J., Guillory, J., Durinck, S., Vilgalys, T. P., Kwok, P.-Y., Cox, L. A., Seshagiri, S., Song, Y. S., & Wall, J. D. (2020). Accurate assembly of the olive baboon ( *Papio anubis* ) genome using long-read and Hi-C data. *GigaScience*, *9*(12), giaa134. https://doi.org/10.1093/gigascience/giaa134

Boyle, P., Clement, K., Gu, H., Smith, Z. D., Ziller, M., Fostel, J. L., Holmes, L., Meldrim, J., Kelley, F., Gnirke, A., & Meissner, A. (2012). Gel-free multiplexed reduced representation bisulfite sequencing for large-scale DNA methylation profiling. *Genome Biology*, *13*(10), R92. https://doi.org/10.1186/gb-2012-13-10-r92

Fan, Y., Vilgalys, T. P., Sun, S., Peng, Q., Tung, J., & Zhou, X. (2019). IMAGE: High-powered detection of genetic effects on DNA methylation using integrated methylation QTL mapping and allele-specific analysis. *Genome Biology*, *20*(1), 220. https://doi.org/10.1186/s13059-019-1813-1

Guo, W., Zhu, P., Pellegrini, M., Zhang, M. Q., Wang, X., & Ni, Z. (2018). CGmapTools improves the precision of heterozygous SNV calls and supports allele-specific methylation detection and visualization in bisulfite-sequencing data. *Bioinformatics*, *34*(3), 381–387. https://doi.org/10.1093/bioinformatics/btx595

Krueger, F., & Andrews, S. R. (2011). Bismark: A flexible aligner and methylation caller for Bisulfite-Seq applications. *Bioinformatics*, *27*(11), 1571–1572. https://doi.org/10.1093/bioinformatics/btr167

Langmead, B., & Salzberg, S. L. (2012). Fast gapped-read alignment with Bowtie 2. *Nature Methods*, *9*(4), 357–359. https://doi.org/10.1038/nmeth.1923

Lea, A. J., Altmann, J., Alberts, S. C., & Tung, J. (2016). Resource base influences genome-wide DNA methylation levels in wild baboons (Papio cynocephalus). *Molecular Ecology*, *25*(8), 1681–1696. https://doi.org/10.1111/mec.13436

Lea, A. J., Tung, J., & Zhou, X. (2015). A Flexible, Efficient Binomial Mixed Model for Identifying Differential DNA Methylation in Bisulfite Sequencing Data. *PLOS Genetics*, *11*(11), e1005650. https://doi.org/10.1371/journal.pgen.1005650

Lea, A. J., Vockley, C. M., Johnston, R. A., Del Carpio, C. A., Barreiro, L. B., Reddy, T. E., & Tung, J. (2018). Genome-wide quantification of the effects of DNA methylation on human gene regulation. *eLife*, *7*, e37513. https://doi.org/10.7554/eLife.37513

Leek, J. T., Johnson, W. E., Parker, H. S., Jaffe, A. E., & Storey, J. D. (2012). The sva package for removing batch effects and other unwanted variation in high-throughput experiments. *Bioinformatics*, *28*(6), 882–883. https://doi.org/10.1093/bioinformatics/bts034

McKenna, A., Hanna, M., Banks, E., Sivachenko, A., Cibulskis, K., Kernytsky, A., Garimella, K., Altshuler, D., Gabriel, S., Daly, M., & DePristo, M. A. (2010). The Genome Analysis Toolkit: A MapReduce framework for analyzing next-generation DNA sequencing data. *Genome Research*, *20*(9), 1297–1303. https://doi.org/10.1101/gr.107524.110

Muggeo, V. M. R. (2016). Testing with a nuisance parameter present only under the alternative: A score-based approach with application to segmented modelling. *Journal of Statistical Computation and Simulation*, *86*(15), 3059–3067. https://doi.org/10.1080/00949655.2016.1149855

Robinson, J. A., Belsare, S., Birnbaum, S., Newman, D. E., Chan, J., Glenn, J. P., Ferguson, B., Cox, L. A., & Wall, J. D. (2019). Analysis of 100 high-coverage genomes from a pedigreed captive baboon colony. *Genome Research*, *29*(5), 848–856. https://doi.org/10.1101/gr.247122.118

Hastie, R. T. (2017). *Impute* [Computer software]. Bioconductor. https://doi.org/10.18129/B9.BIOC.IMPUTE

Tung, J., Primus, A., Bouley, A. J., Severson, T. F., Alberts, S. C., & Wray, G. A. (2009). Evolution of a malaria resistance gene in wild primates. *Nature*, *460*(7253), 388–391. https://doi.org/10.1038/nature08149

Tung, J., Zhou, X., Alberts, S. C., Stephens, M., & Gilad, Y. (2015). The genetic architecture of gene expression levels in wild baboons. *eLife*, *4*, e04729. https://doi.org/10.7554/eLife.04729

Vilgalys, T. P., Fogel, A. S., Anderson, J. A., Mututua, R. S., Warutere, J. K., Siodi, I. L., Kim, S. Y., Voyles, T. N., Robinson, J. A., Wall, J. D., Archie, E. A., Alberts, S. C., & Tung, J. (2022). Selection against admixture and gene regulatory divergence in a long-term primate field study. *Science*, *377*(6606), 635–641. https://doi.org/10.1126/science.abm4917

Vilgalys, T. P., Rogers, J., Jolly, C. J., Baboon Genome Analysis, Mukherjee, S., & Tung, J. (2019). Evolution of DNA Methylation in Papio Baboons. *Molecular Biology and Evolution*, *36*(3), 527–540. https://doi.org/10.1093/molbev/msy227

Wall, J. D., Schlebusch, S. A., Alberts, S. C., Cox, L. A., Snyder-Mackler, N., Nevonen, K. A., Carbone, L., & Tung, J. (2016). Genomewide ancestry and divergence patterns from low-coverage sequencing data reveal a complex history of admixture in wild baboons. *Molecular Ecology*, *25*(14), 3469–3483. https://doi.org/10.1111/mec.13684

Xi, Y., & Li, W. (2009). BSMAP: Whole genome bisulfite sequence MAPping program. *BMC Bioinformatics*, *10*(1), 232. https://doi.org/10.1186/1471-2105-10-232

### Supplementary Figures

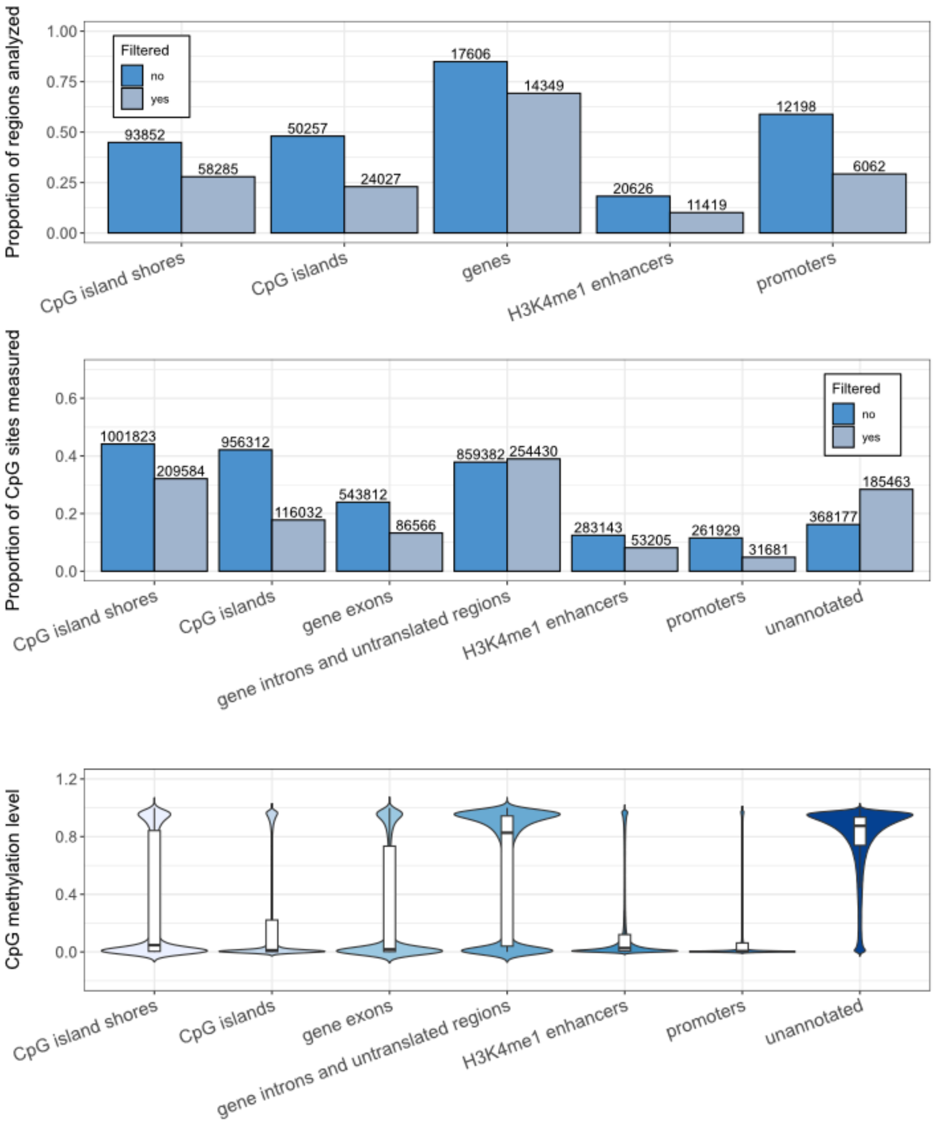


**Figure S1: Properties of the Amboseli baboon RRBS data set. (A)** Proportion of annotated features in the baboon genome for which at least one CpG site was analyzed. Numbers above each bar represent the number of features tagged in our data set. **(B)** Proportion of total CpG sites analyzed that fell in each genomic region. Numbers above each bar represent the number of CpG sites included from each category (note that the same CpG site can fall in more than one category). In (A) and (B), dark blue bars show all sites with 5x coverage (n = 2.2 million sites); light blue bars show sites filtered for mean methylation between 10% and 90% (n = 653k sites). **(C)** Violin plots showing the distribution of mean DNA methylation levels for CpG sites located in each genomic context, before filtering. White box plots show the interquartile range and median (black bar).

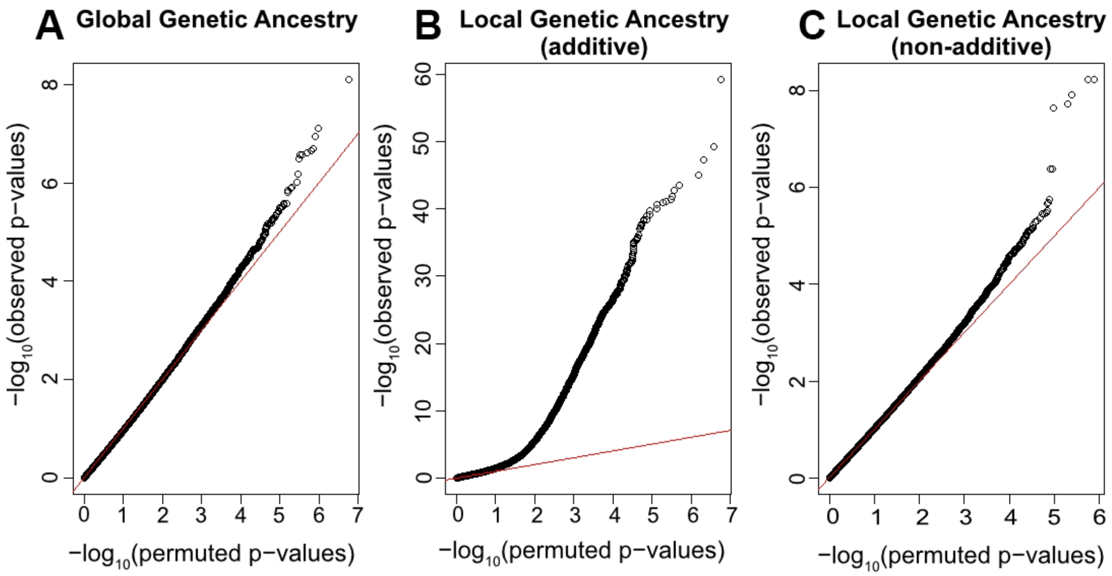


**Figure S2: Signal for ancestry effects on DNA methylation in Amboseli.** Quantile-quantile plots that show the enrichment of associations between DNA methylation and **(A)** global genetic ancestry; **(B)** local ancestry, modeled additively; and **(C)** local ancestry, modeled non-additively. In each case, the y-axis represents the observed distribution of $-\log_{10}$ p-values from a mixed effects model with DNA methylation as the response variable and global/local ancestry as the predictor of interest. The x-axis represents the distribution of $-\log_{10}$ p-values obtained from 10 permutations of the data (see Methods), and the red x=y line shows the expectation for a Q-Q plot under a scenario of no enrichment.

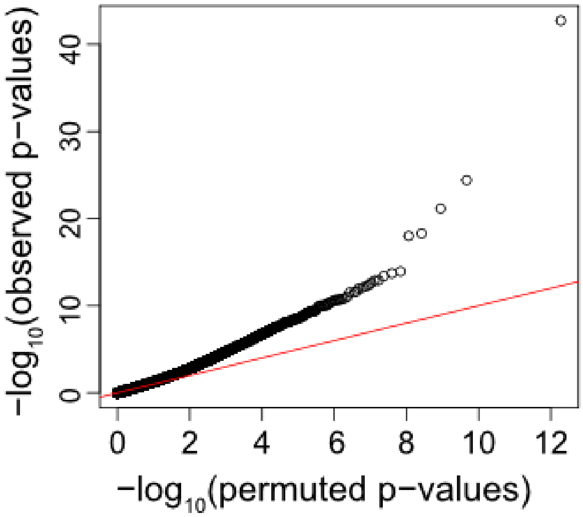


**Figure S3: Differences in DNA methylation between anubis and yellow baboons, sampled outside of the hybrid zone.** Quantile-quantile plots that show the enrichment of associations between DNA methylation and species identity (yellow vs. anubis baboons). The y-axis represents the observed distribution of $-\log_{10}$ p-values with DNA methylation as the response variable and species as the predictor of interest. The x-axis represents the distribution of $-\log_{10}$ p-values obtained from 10 permutations of the data (see Methods), and the red x=y line shows the expectation for a Q-Q plot under a scenario of no enrichment.

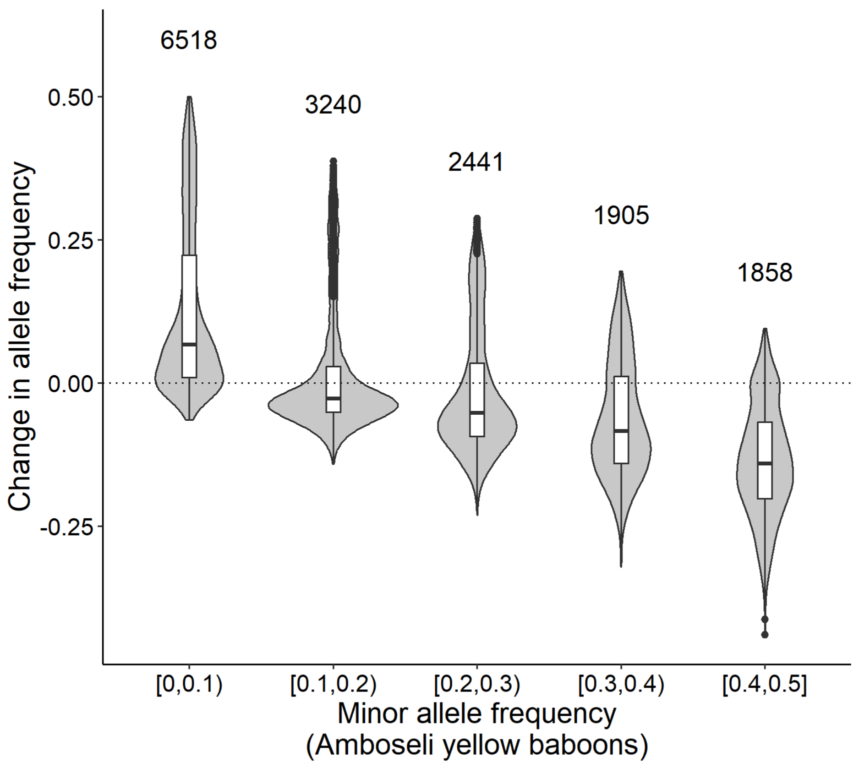


**Figure S4: Change in allele frequency depends on the unadmixed minor allele frequency.** Violin plots showing the change in minor allele frequency at meQTL loci, binned by the minor allele frequency in unadmixed yellow baboons. Numbers above each violin plot represent the number of CpG sites in that bin. As in Fig 3A, minor allele frequencies were calculated for all individuals with locally homozygous yellow baboon ancestry. These values were compared against those for all genotyped individuals in the Amboseli population. The shift towards higher minor allele frequencies is driven by variants which are relatively rare in yellow baboons (i.e., the first bin).

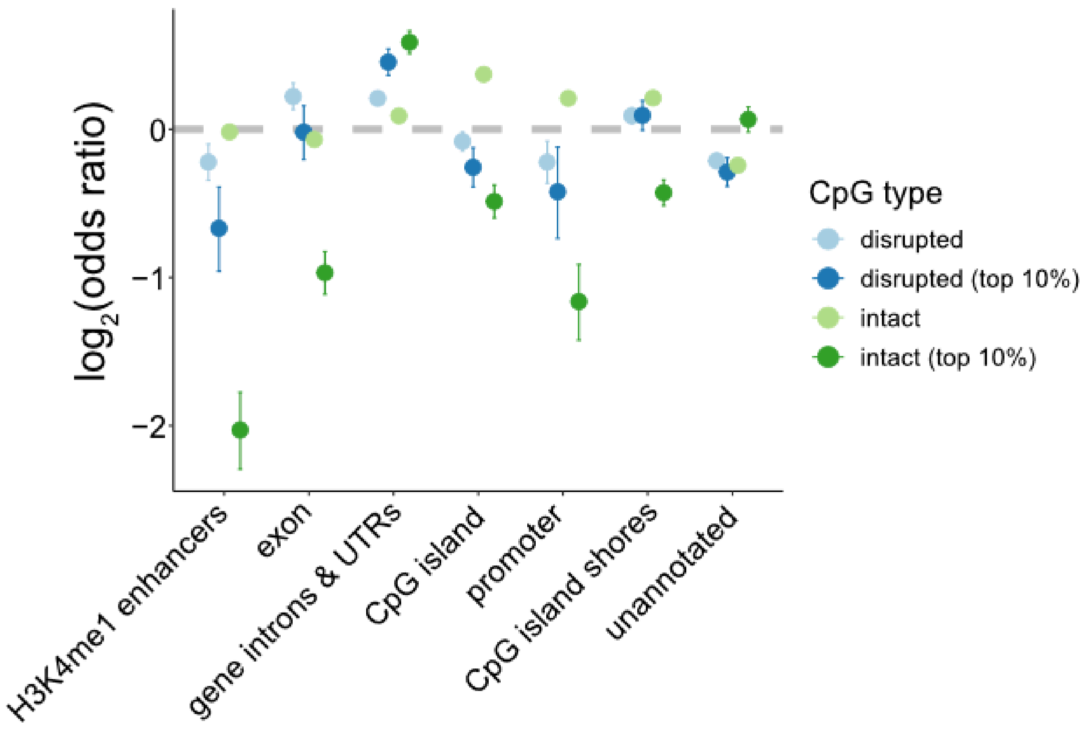


**Figure S5: Enrichment of ancestry-associated methylation by genomic context.** Ancestry-associated CpG sites are stratified into those where the CpG site is not disrupted by a SNP (green) and those that are disrupted by CpG site-abolishing SNP (blue), as these sites show different patterns of enrichment. In both sets, enrichment statistics are magnified for the top 10% of local ancestry effects. This comparison makes the effect sizes between intact and disrupted sites more congruent, and also makes the pattern of enrichment by genomic context more congruent.

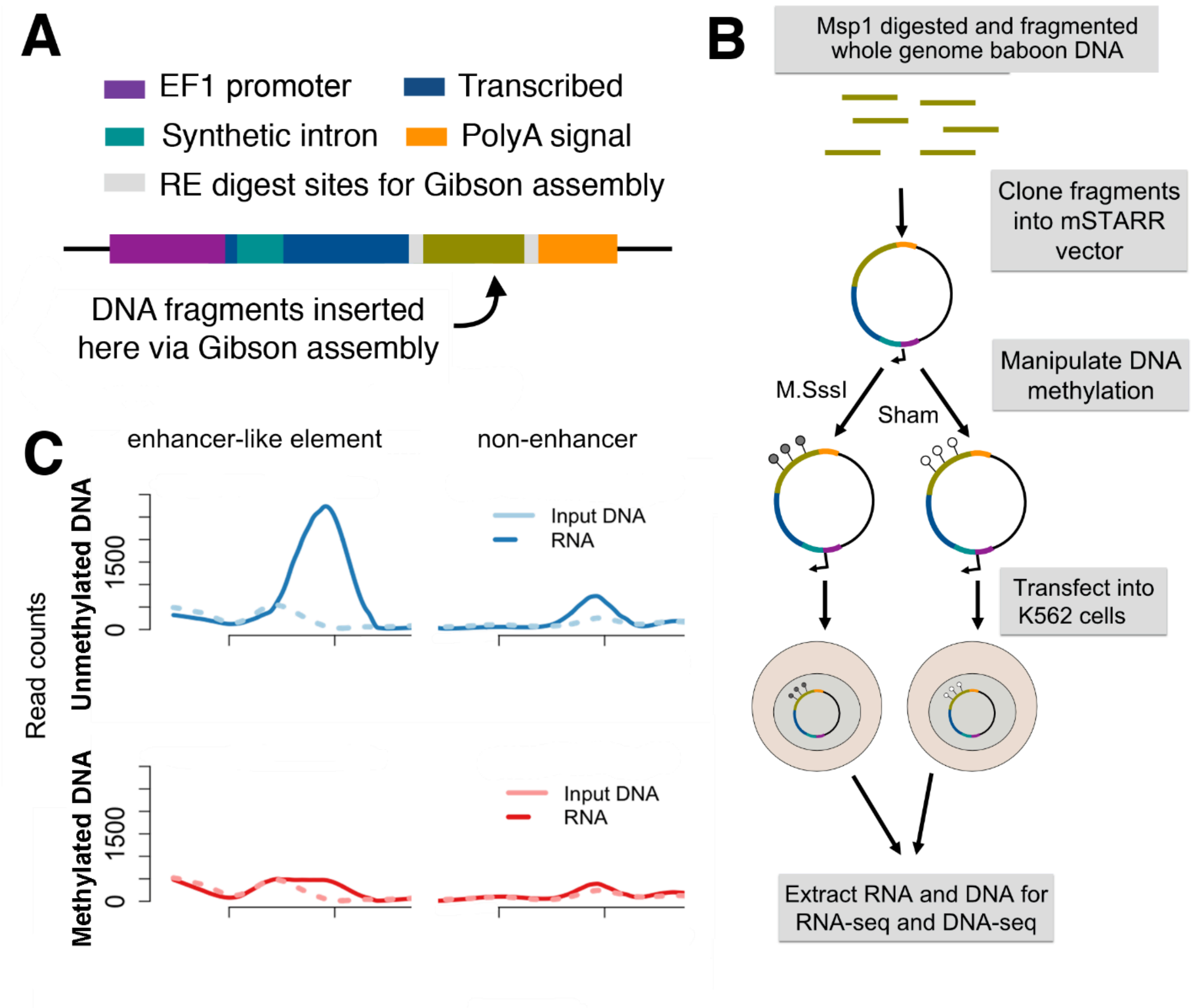


**Figure S6: mSTARR-seq assay. (A)** The CpG *pmSTARRseq* vector is designed so that functional regulatory elements will self-transcribe to produce an mRNA transcript, including a transcribed region (dark blue) that spans a synthetic intron (teal), the sequence of the regulatory element itself (olive green), and an SV40 polyA signal (orange). **(B)** Input DNA fragments from an anubis baboon were cloned into the mSTARR-seq plasmid vector and subjected to experimental methylation (*M.Sss1* treatment), which methylates CpG motifs, or a sham treatment (buffer only), which leaves them unmethylated. Plasmids were then transfected into K562 cells with 6 technical replicates per condition. Plasmid DNA and plasmid-derived mRNA was then extracted and the variable insert regions sequenced. **(C)** An example region where regulatory activity depends upon DNA methylation, and a second example in which there is no enhancer activity. More RNA-seq reads than DNA-seq reads identify genomic regions with regulatory activity. (Figure modified from Lea et al., 2018).
